# A systematic comparison of chloroplast genome assembly tools

**DOI:** 10.1101/665869

**Authors:** Jan A Freudenthal, Simon Pfaff, Niklas Terhoeven, Arthur Korte, Markus J Ankenbrand, Frank Förster

## Abstract

**Background:** Chloroplasts are intracellular organelles that enable plants to conduct photosynthesis. They arose through the symbiotic integration of a prokaryotic cell into an eukaryotic host cell and still contain their own genomes with distinct genomic information. Plastid genomes accommodate essential genes and are regularly utilized in biotechnology or phylogenetics. Different assemblers that are able to assess the plastid genome have been developed. These assemblers often use data of whole genome sequencing experiments, which usually contain reads from the complete chloroplast genome.

**Results:** The performance of different assembly tools has never been systematically compared. Here we present a benchmark of seven chloroplast assembly tools, capable of succeeding in more than 60% of known real data sets. Our results show significant differences between the tested assemblers in terms of generating whole chloroplast genome sequences and computational requirements. The examination of 105 data sets from species with unknown plastid genomes leads to the assembly of 20 novel chloroplast genomes.

**Conclusions:** We create docker images for each tested tool that are freely available for the scientific community and ensure reproducibility of the analyses. These containers allow the analysis and screening of data sets for chloroplast genomes using standard computational infrastructure. Thus, large scale screening for chloroplasts within genomic sequencing data is feasible.

## Introduction

### General introduction and motivation

Chloroplasts are essential organelles present in the cells of plants and autotrophic protists, which enable the conversion of light energy into chemical energy via photosynthesis. They harbor their own prokaryotic type of ribosomes and a circular DNA genome that varies in size between 120 kbp to 160 kbp [1]. Because of their small size, chloroplast genomes were one of the first targets for sequencing projects. The first chloroplast genome sequences were obtained in 1986 [2, 3]. These early efforts elucidated the general genome organization and structure of the chloroplast DNA and have been reviewed previously [4, 5]. Chloroplast genomes are widely used for evolutionary analyses [6, 7], barcoding [8, 9, 10], and meta-barcoding [11, 12]. Interesting features of chloroplast genomes include their small size (120 kbp to 160 kbp,[1]), due to endosymbiotic gene transfer [13, 14], and the low number of 100 to 120 genes that are encoded within the genome [4]. Despite the overall high sequence conservation of the chloroplast genome, there are striking differences in the gene content between different autotroph groups, exemplified by the loss of the whole *ndh* gene family in Droseraceae [15]). Even more extreme evolutionary cases, where chloroplasts show a very low GC content and a modified genetic code have been described [16].

Structurally, two inverted repeats (Inverted Repeats (IRs)) named IR_A_ and IR_B_ of 10 kbp to 76 kbp divide the chloroplast genome into a Large Single Copy (LSC) and a Small Single Copy (SSC) region [1], which complicates genome assembly with short read technologies[17]. Moreover, the existence of different chloroplasts within a single individual, and thus multiple different chloroplast genomes, have been described for various plants [18, 19, 20]. This phenomenon - called heteroplasmy - is only poorly understood in terms of its origin and evolutionary importance, but it impacts the assembly of whole chloroplast genomes.

Nonetheless, given its small size, it is still much easier to decipher a complete chloroplast genome than a complete core genome. Consequently, many comparative genomic approaches target the chloroplast genome. For example the *Arabidopsis thaliana* core genome is approximately 125 Mbp in length [21, 22] while the size of the *A. thaliana* chloroplast genome at 154 kbp is more than 800 times smaller [23].

Each single chloroplast contains several hundred copies of its genome [24, 25]. Therefore, many plant core genome sequencing projects contain reads that originate from chloroplasts as a by-product and permit the assembly of chloroplast genomes. Such sequences are available from databases such as the Sequence Read Archive at NCBI [26].

Complete chloroplast genomes can be used as super-barcodes [27], both for biotechnology applications and genetic engineering [28]. Furthermore, the availability of whole chloroplast genomes would enable large scale comparative studies [29].

### Approaches to extracting chloroplasts sequences from whole genome data

Different strategies have been developed to assemble chloroplast genomes [30]. In general, obtaining a chloroplast genome from whole genome sequencing (WGS) data requires two steps: (1) extraction of chloroplast reads from the sequencing data; (2) assembly and resolution of the special circular structure including the IRs. The extraction of chloroplast reads can be achieved by mapping the reads to a reference chloroplast. [31]. A different approach that does not depend on the availability of a reference chloroplast, uses the higher coverage of reads originating from the chloroplast [32]. Here, a *k*-mer analysis can be used to extract the most frequent reads. An example for this is implemented in `chloroExtractor` [33]. A third method, which is for example used by `NOVOPlasty` [34], combines both approaches by using a reference chloroplast as seed and simultaneously assembling the reads based on *k*-mers.

### Purpose and scope of this study

The goal of this study was to compare the effectiveness and efficiency of existing open source command-line tools to perform a de-novo assembly of whole chloroplast genomes from raw genomic data. We only compared tools that require minimal configuration, which includes no need for extensive data preparation, no need for a specific reference (apart from *A. thaliana*), no need to change default parameters, and no manual finishing. We further restricted our benchmark to paired end Illumina data as the sole input, as these are routinely generated by modern sequencing platforms [35].

Thus our analyses reflect the most common use cases: (1) trying a tool quickly without digging into options for fine tuning; (2) large scale automatic applications. We tested all tools on more than 100 real data sets for species without published chloroplast genomes. The performance of most tools might be significantly improved by optimizing parameters for each data set specifically, but this exhaustive comparison - including tuning of all different possible parameters for each tool-was out of the scope of this study.

To summarize, we provide new chloroplast genome sequences for many species and demonstrated the ability to discover and assemble novel chloroplast genomes as well as asses inter/intra-individual differences in the respective chloroplast genomes.

## Results

### Performance metrics

All described tools have been tested with regard to their assembly time, memory and CPU utilization.

#### Time requirements

Massive differences between the different tools were observed in terms of the run time for the assembly. Apart from tool-specific differences, input data and number of threads used had a huge impact on the time requirement. The observed run times varied from a few minutes to several hours (Figure 1).

**Figure 1.**
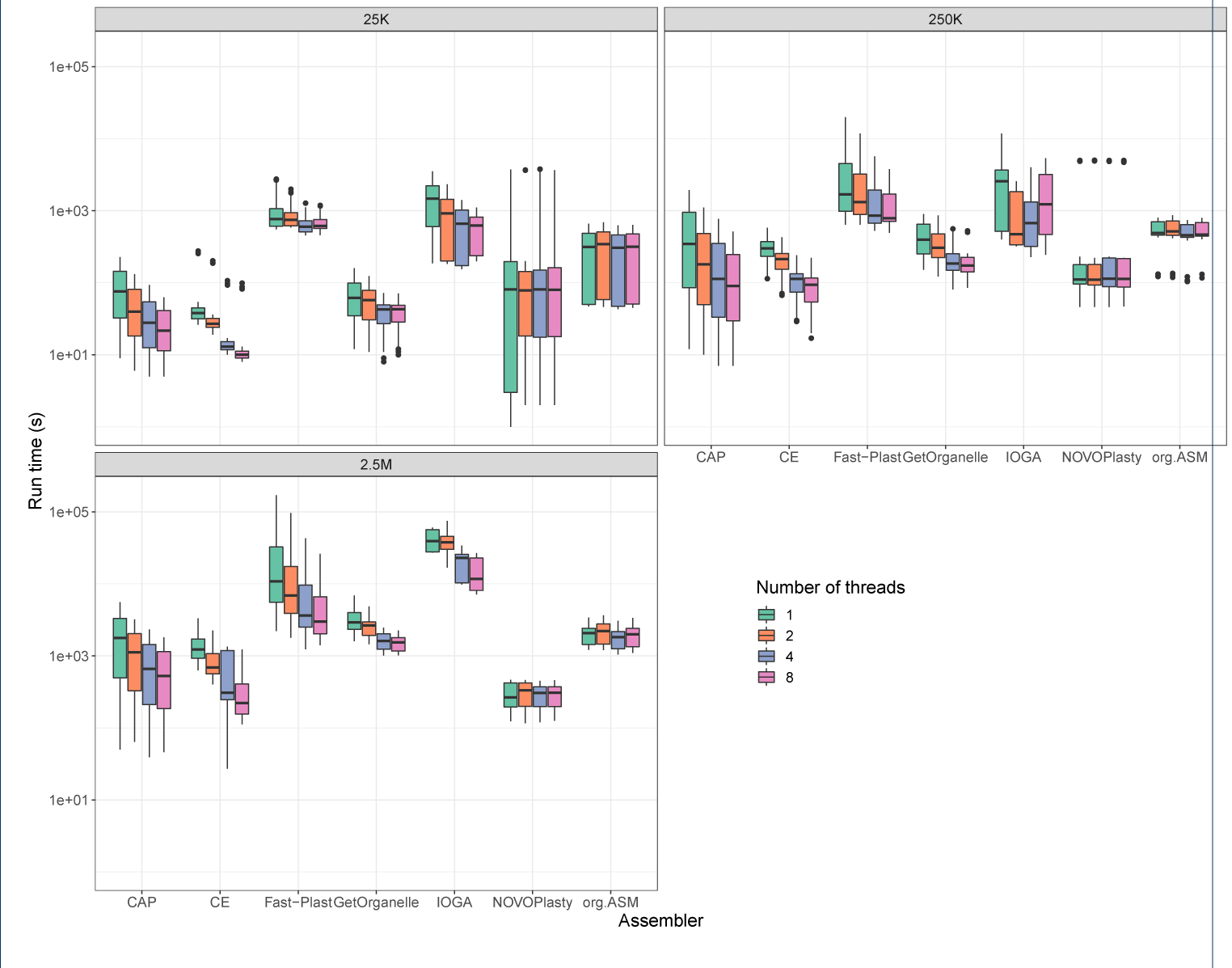
Computation time depending on number of threads and size of input data. The boxplots show the differences in demand of CPU time for different number of threads and input data size for the seven different assemblers

Some assemblies failed to finish within our time limit of 48 h. On average, the longest time to generate an assembly was taken by `IOGA` and `Fast-Plast` followed by `ORG.Asm` and `GetOrganelle`. The most time efficient tool was `chloroExtractor`, which was a little faster than `NOVOPlasty` and `Chloroplast assembly protocol`. Not all tested tools were able to benefit from having access to multiple threads. Both `NOVOPlasty` and `ORG.Asm` required almost the same time independent of being allowed to utilize 1, 2, 4, or 8 threads. In contrast, `Chloroplast assembly protocol`, `chloroExtractor`, `GetOrganelle`and `Fast-Plast` all profited from multi-threading settings (Figures 1 and 2 and Additional file 1: Tables S4 to S6).

**Figure 2.**
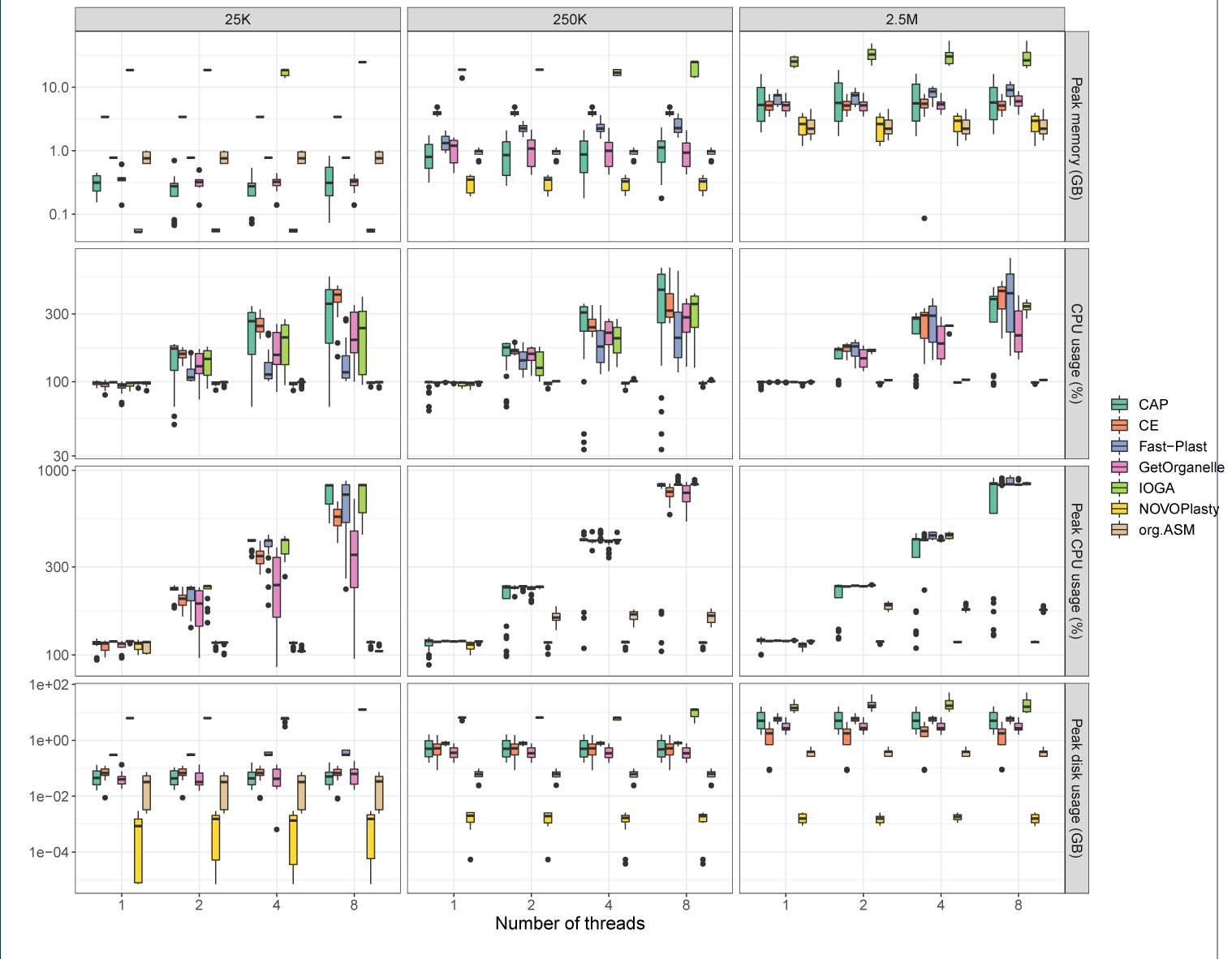
Performance metrics. Boxplots depicting the demand of CPU and RAM and disk space needed depending on the assembler, input data size and number of threads

#### Memory and CPU Usage

The peak and mean CPU usage, as well as peak memory and disk usage were recorded for all assemblers based on the same input data set and number of threads (Figure 2 and Additional file 1: Tables S4 to S6). In general, the size of the input data influenced the peak memory usage with the exception of `chloroExtractor` and `IOGA`. Those two assemblers showed a memory usage pattern, which was less influenced by the size of the data. The number of allowed threads had only a limited impact on the peak memory usage. All programs profited from a higher number of threads, if the size of the input data was increased concerning their memory and CPU usage footprint. In contrast, the disk usage was independent of the size of the input data and the number of threads for all assemblers.

### Qualitative

On average, the user experience in terms of installation and running the analyses was evaluated as Good for all tools (Table 1).

**Table 1.** Overview of the results of the qualitative usability evaluation. Each tool could score Good, Okay or Bad in each of the categories.

| Tool | Installation | Test/Tutorial | Documentation | Maintenance | FLOSS |
| --- | --- | --- | --- | --- | --- |
| chloroExtractor | GOOD | GOOD | GOOD | GOOD | GOOD |
| Chloroplast assembly protocol | OKAY | GOOD | OKAY | GOOD | GOOD |
| Fast-Plast | BAD | OKAY | GOOD | GOOD | GOOD |
| GetOrganelle | OKAY | OKAY | GOOD | GOOD | GOOD |
| IOGA | BAD | BAD | OKAY | BAD | GOOD |
| NOVOPlasty | GOOD | GOOD | GOOD | GOOD | OKAY |
| ORG.Asm | BAD | BAD | OKAY | GOOD | GOOD |

**Table 2.**
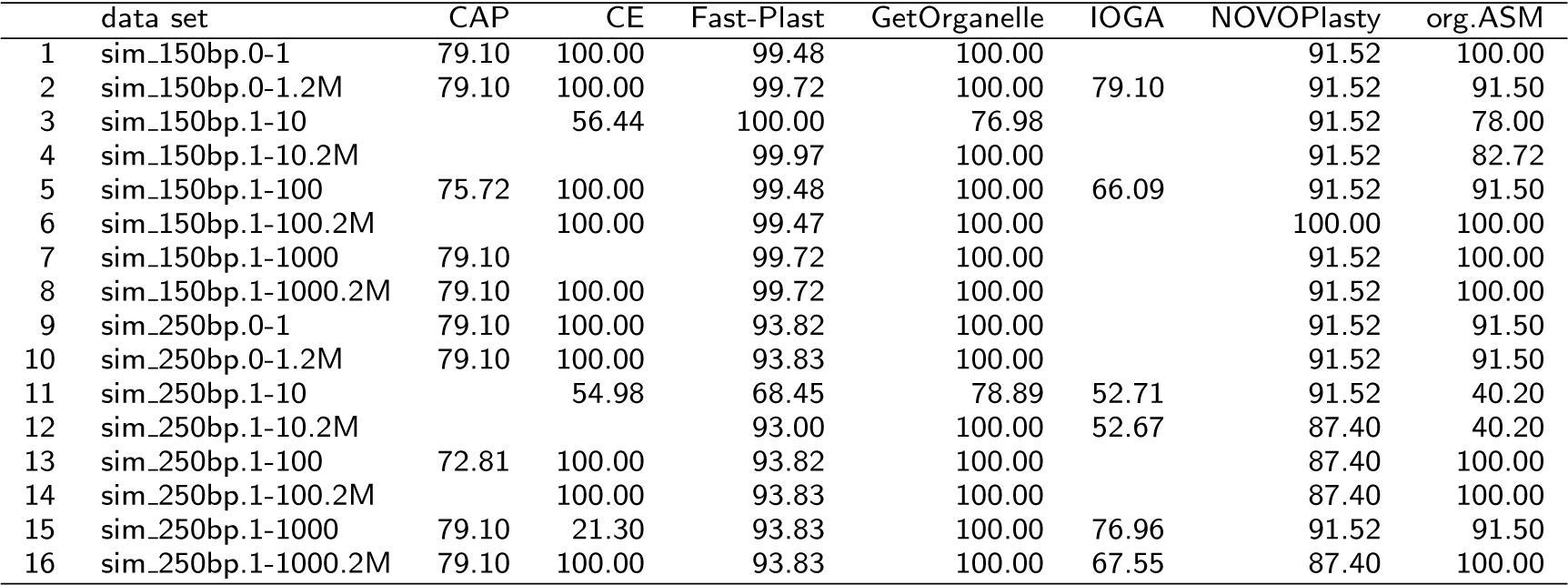
Scores of assemblies of simulated data

|  | data set | CAP | CE | Fast-Plast | GetOrganelle | IOGA | NOVOPlasty | org.ASM |
| --- | --- | --- | --- | --- | --- | --- | --- | --- |
| 1 | sim_150bp.0-1 | 79.10 | 100.00 | 99.48 | 100.00 |  | 91.52 | 100.00 |
| 2 | sim_150bp.0-1.2M | 79.10 | 100.00 | 99.72 | 100.00 | 79.10 | 91.52 | 91.50 |
| 3 | sim_150bp.1-10 |  | 56.44 | 100.00 | 76.98 |  | 91.52 | 78.00 |
| 4 | sim_150bp.1-10.2M |  |  | 99.97 | 100.00 |  | 91.52 | 82.72 |
| 5 | sim_150bp.1-100 | 75.72 | 100.00 | 99.48 | 100.00 | 66.09 | 91.52 | 91.50 |
| 6 | sim_150bp.1-100.2M |  | 100.00 | 99.47 | 100.00 |  | 100.00 | 100.00 |
| 7 | sim_150bp.1-1000 | 79.10 |  | 99.72 | 100.00 |  | 91.52 | 100.00 |
| 8 | sim_150bp.1-1000.2M | 79.10 | 100.00 | 99.72 | 100.00 |  | 91.52 | 100.00 |
| 9 | sim_250bp.0-1 | 79.10 | 100.00 | 93.82 | 100.00 |  | 91.52 | 91.50 |
| 10 | sim_250bp.0-1.2M | 79.10 | 100.00 | 93.83 | 100.00 |  | 91.52 | 91.50 |
| 11 | sim_250bp.1-10 |  | 54.98 | 68.45 | 78.89 | 52.71 | 91.52 | 40.20 |
| 12 | sim_250bp.1-10.2M |  |  | 93.00 | 100.00 | 52.67 | 87.40 | 40.20 |
| 13 | sim_250bp.1-100 | 72.81 | 100.00 | 93.82 | 100.00 |  | 87.40 | 100.00 |
| 14 | sim_250bp.1-100.2M |  | 100.00 | 93.83 | 100.00 |  | 87.40 | 100.00 |
| 15 | sim_250bp.1-1000 | 79.10 | 21.30 | 93.83 | 100.00 | 76.96 | 91.52 | 91.50 |
| 16 | sim_250bp.1-1000.2M | 79.10 | 100.00 | 93.83 | 100.00 | 67.55 | 87.40 | 100.00 |

However, we discovered the following slight problems:

Two minor dependencies were missing in the `GetOrganelle` installation instructions and there were no test data available [36]. Additionally, an issue occurred when running it on one particular *A. thaliana* data set. This was resolved after contact with the authors via GitHub [37].

The `Fast-Plast` installation instructions were missing some dependencies [38]. Like `GetOrganelle`, `Fast-Plast` does not offer a test data set or a tutorial, except for some example commands [36].

The `ORG.Asm` installation instructions did not work. We found some issues, which were probably related to the requirement of `Python 3.7` [39]. A tutorial including sample data was available, but following the instructions resulted in a segmentation fault. We found a workaround for this bug and contacted the authors [40].

The main critique point of `NOVOPlasty` was the lack of test data and instructions. This was fixed by the authors after we contacted them [41]. Additionally, `NOVOPlasty` uses a custom license, where an OSI approved license would be preferable.

The `chloroExtractor` does come with test data and a short tutorial. However, it is currently not possible to evaluate the results of the test run as the expected results are not available [42].

The `IOGA` installation instructions were missing many dependencies [43]. There was also no test data or tutorial available and no license assigned to it [44]. After contacting the authors, the AGPL-3.0 license was added [45], as well as a note in the description explaining, that `IOGA` is no longer maintained.

Installation instructions for `Chloroplast assembly protocol` were also missing some dependencies. The list was updated after we contacted the authors [46]. This tool does come with an extensive tutorial and test data, but the expected outcome is not provided.

### Quantitative

For a quantitative evaluation we tested the capacity of all programs to assemble chloroplasts based on different input data. Input data were either generated from existing chloroplast genomes or downloaded from sequencing repositories.

#### Simulated data

The different simulated data were all based on the *A. thaliana* chloroplast and core genome sequence. Some general trends could be observed: a ratio of 1:10 genome to chloroplast reads, contains too few chloroplastic reads for most tools (except `Fast-Plast` and `GetOrganelle`). A good performance for all tools was observed at a ratio of 1:100. Increasing the ratio further had no additional benefit, even if pure chloroplast reads were used (Figure 3). Using 250 bp paired read compared to 150 bp paired reads, did not produce improved results (Figure 3). In the case of `Fast-Plast`, the performance was even worse with the longer read length as more than a single copy of the chloroplast genome was returned.

**Figure 3.**
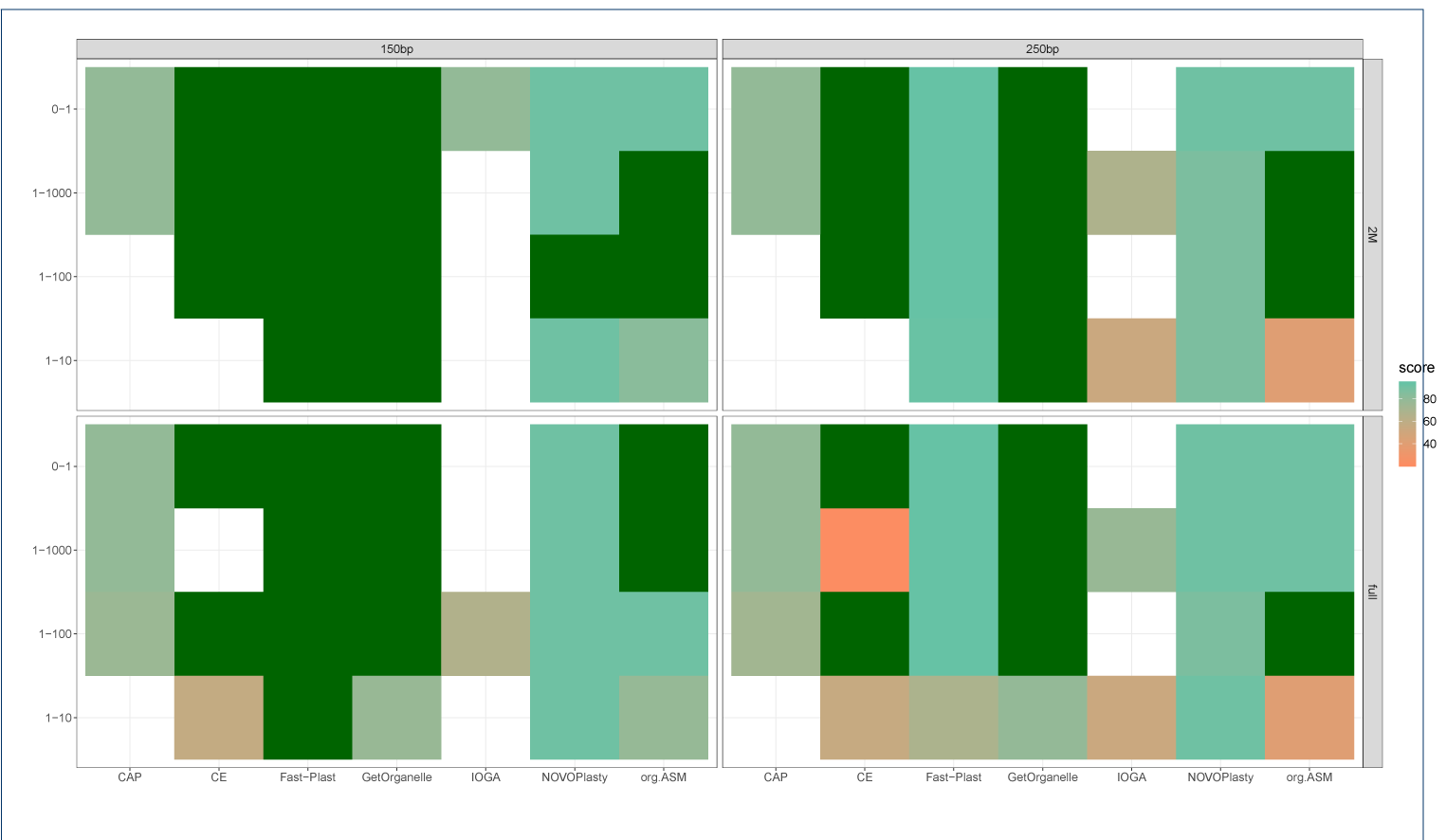
Score of assemblies on simulated data. Results of assemblies from simulated data sets. Color scale of the tiles represents the score

Overall `GetOrganelle` and `Fast-Plast` were the most successful tools on the simulated data while `Chloroplast assembly protocol` and `IOGA` were unable to successfully assemble any chloroplasts out of the 16 different data sets.

#### Real data sets

To evaluate the performance on real data, we used publicly available short read data from NCBI’s SRA with existing reference chloroplasts. We observed considerable differences for the tested assemblers, if we compared the generated alignments against the reference chloroplasts (Figure 4). The highest scores were achieved by `GetOrganelle` with a median of 99.8 and 210 circular assemblies out of a total of 360 assemblies that resulted in an output (Table 3). The performance of `GetOrganelle` was followed by `Fast-Plast`, `NOVOPlasty`, `IOGA`, and `ORG.Asm`. `Fast-Plast` outperformed `NOVOPlasty` and `ORG.Asm` in terms of score, producing twice as many 113 perfectly assembled chloroplast genomes (`NOVOPlasty` produced 58 and `ORG.Asm` 46 circular genomes). `IOGA` and `Chloroplast assembly protocol` were both unable to assemble a circular, single-contig genome (Table 3, Figure 5).

**Table 3.** Mean scores of chloroplast genome assemblers

|  | assembler | Median | IQR | N_perfect | N_tot |
| --- | --- | --- | --- | --- | --- |
| 1 | CAP | 45.25 | 50.19 | 0 | 369 |
| 2 | CE | 56.55 | 71.50 | 14 | 369 |
| 3 | Fast-Plast | 92.80 | 23.59 | 113 | 369 |
| 4 | GetOrganelle | 99.83 | 20.94 | 210 | 360 |
| 5 | IOGA | 71.10 | 11.21 | 0 | 338 |
| 6 | NOVOPlasty | 75.95 | 48.69 | 58 | 369 |
| 7 | org.ASM | 67.35 | 91.69 | 46 | 348 |

**Figure 4.**
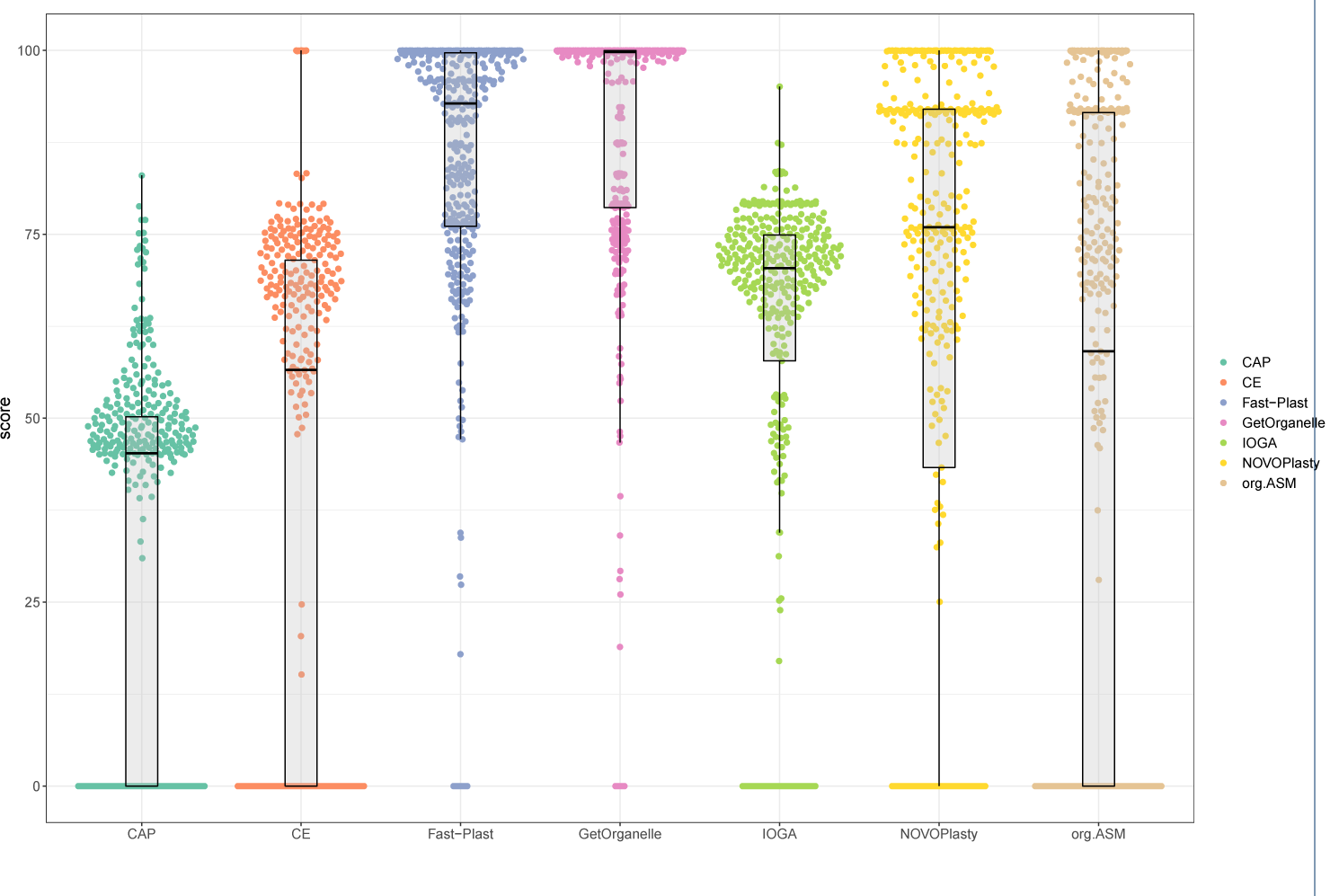
Results of scoring of the seven assemblers. The box- and swarplots depict the results of the scoring algorithm we used. For the different assemblers. The whiskers of boxplots indicate the 1.5 x interquartile range.

**Figure 5.**
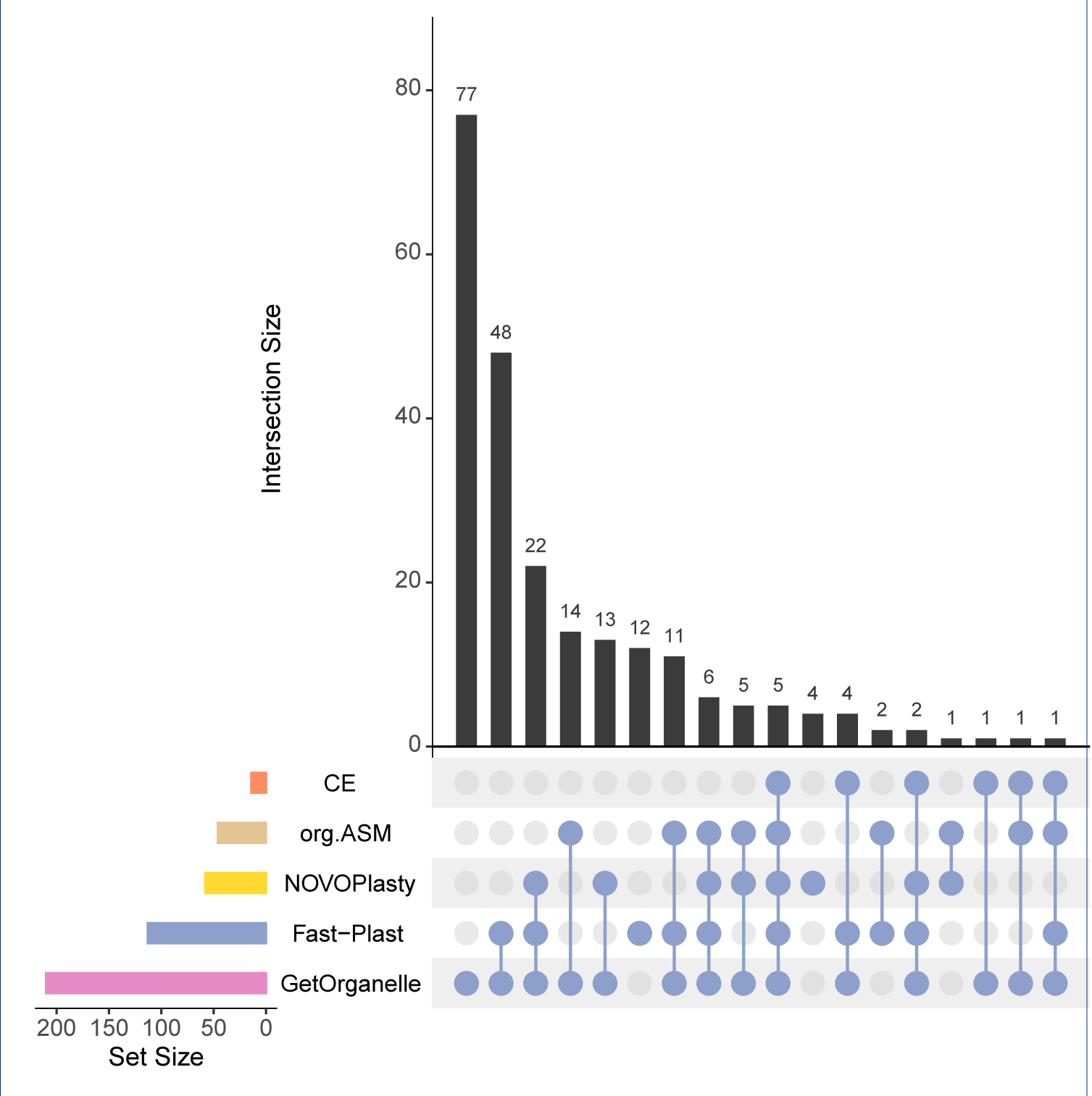
Upset plot [81] comparing success of assemblers on the real data sets. The plot shows the intersection of success (*score* > 99) between assemblers. For 69 data sets only `GetOrganelle` was able to obtain a complete chloroplast. 43 were successful with both `GetOrganelle` and `Fast-Plast` and so on

#### Consistency

Consistency was tested by re-running assemblies using the real data and comparison of the two assemblies (Figure 6). `chloroExtractor` was the only tool able to reproduce the same scores in all runs (Figure 6). `GetOrganelle`, `ORG.Asm`, `Chloroplast assembly protocol`, and `IOGA` generated some assemblies that were unsuccessful in one run, but produced an output in the other attempt. For these assemblers the scores were virtually identical if both runs were succesful. Both `Fast-Plast` and `NOVOPlasty` show only minor changes for the successful assemblies, leading to arrow-shaped scatter plots (Figure 6). `chloroExtractor` appears to be the most robust assembler, showing no deviations between the two runs.

**Figure 6.**
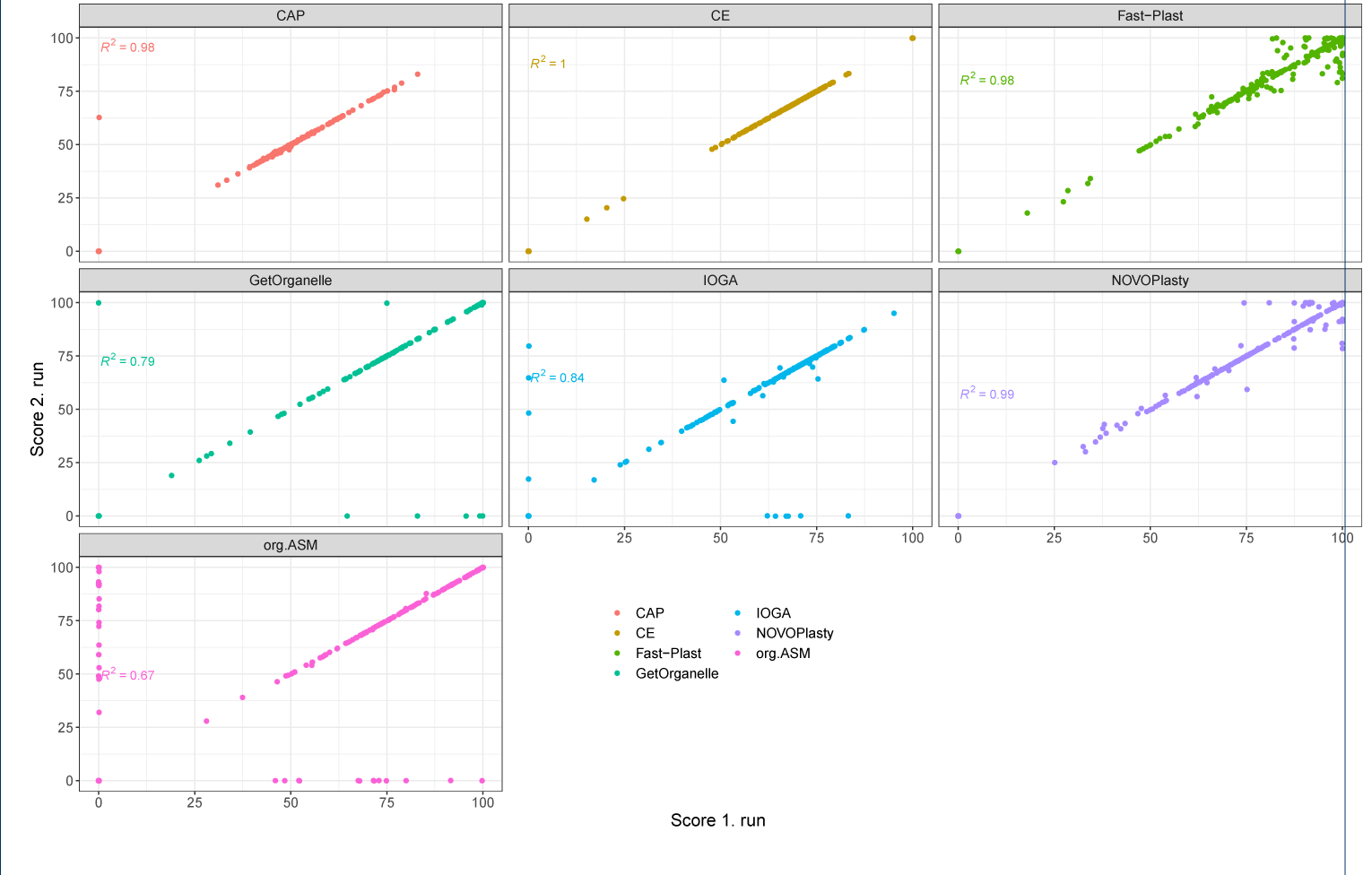
Scores between two repeated runs for consistency testing. The scatter plots depicts the scores of the 1. runs x-axis versus the scores of the 2. run y-axis of the data sets that were selected for re-evaluation.

#### Novel

Finally, the assembly of chloroplasts for species without a published chloroplast, was performed with the different tools. In total 49 out of 105 chloroplasts (46.7 %) with no reference sequence in CpBase were successfully assembled (Figure 8).

**Figure 7.**
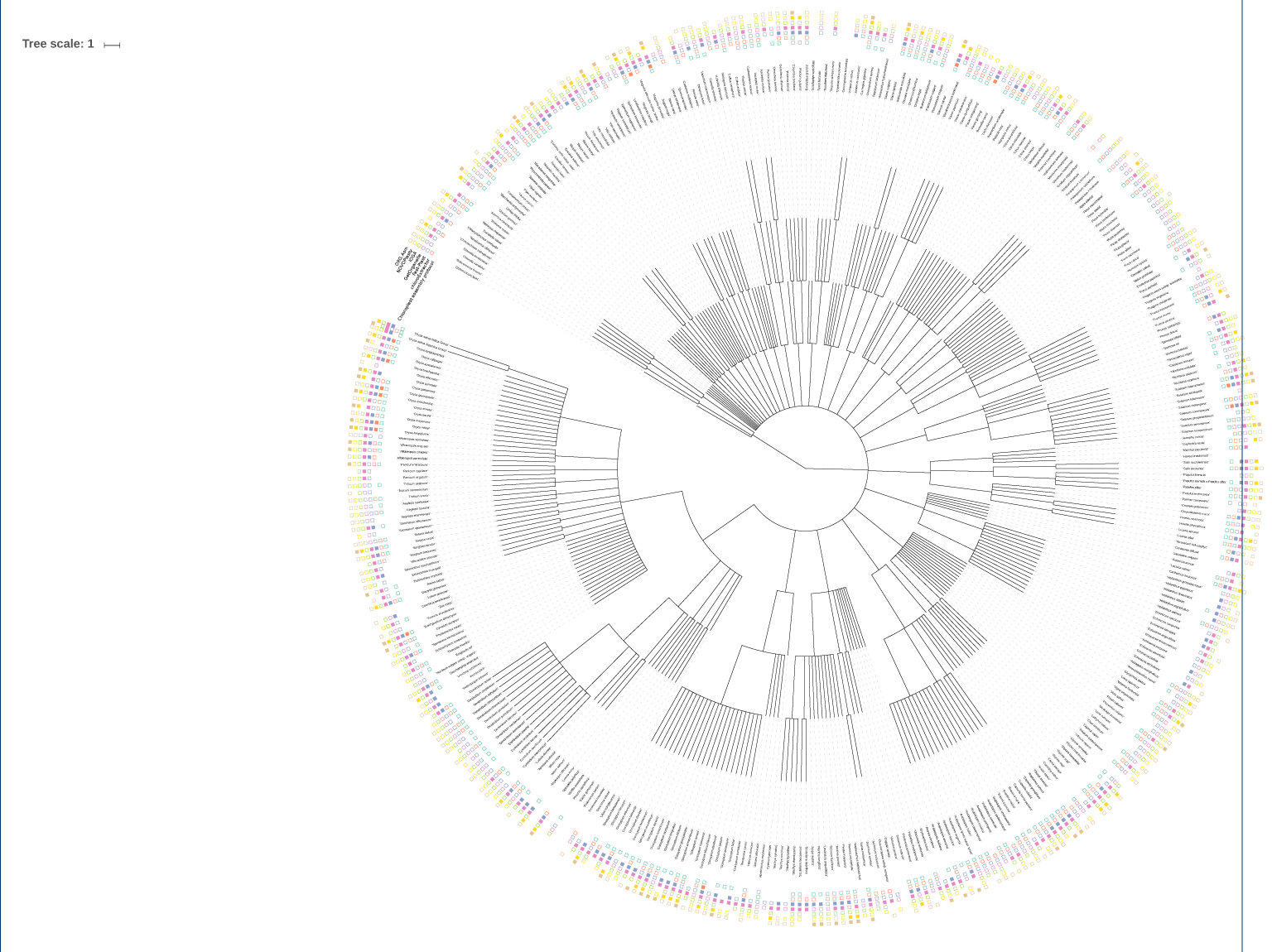
Success for chloroplast assembly shows no taxonomic bias. Success of assemblers on real data sets on tree derived from NCBI taxonomy [82]. Plot was prepared using [83]

**Figure 8.**
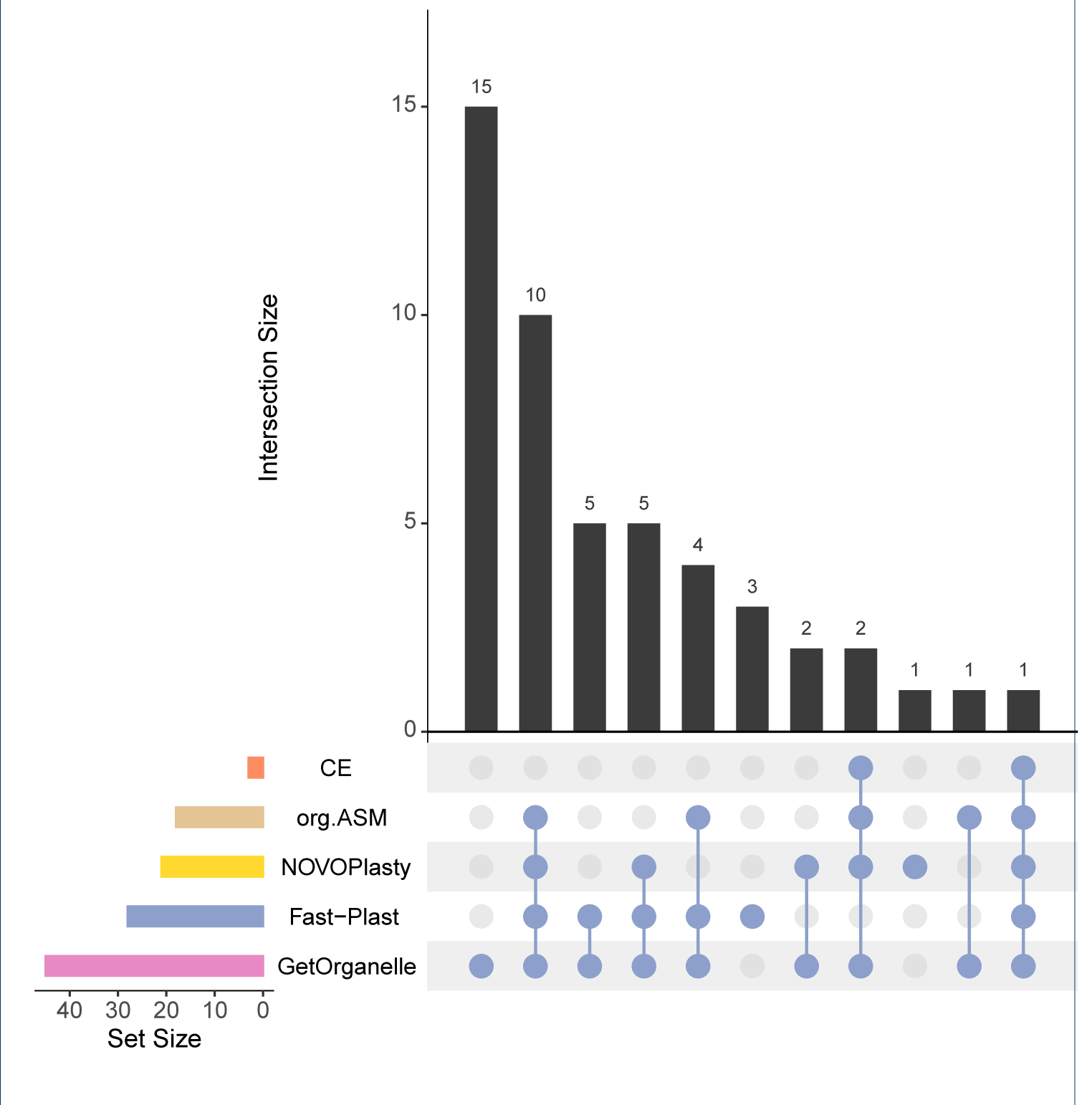
Upset plot [81] comparing success of assemblers on the novel data sets. The plot shows the intersection of success (single contig, *length* ≥ 130*kbp, ir* ≥ 17*kbp*) between assemblers. For 15 data sets only `GetOrganelle` was able to obtain a complete chloroplast. 10 were successful with `GetOrganelle`, `Fast-Plast`, `NOVOPlasty`, and `ORG.Asm` and so on

Almost half (44.9 %) of the successful assembled chloroplasts, were assembled by three or more different tools, while the remaining ones were only successfully generated by one or two different assemblers. Here, `GetOrganelle` showed the best performance and produced 15 distinct chloroplast genomes. For the assemblies obtained from multiple assemblers, we kept the `GetOrganelle` assemblies, after visually inspecting all assemblies using AliTV [47].

For three assemblies, that were obtained by different assemblers, but not by `GetOrganelle`, we kept one assembly obtained by `NOVOPlasty` and two from `Fast-Plast`. All resulting 49 sequences have been annotated with GeSeq [48]. The median number of distinct genes annotated were 80 for coding sequences, 4 for rRNA and 27 for tRNA (Table 5, Figure 10). All sequences were stored in our repository [49]. To avoid multi submissions of the same sequence to Genbank, all 49 sequences have been inspected against Genbank database via BLAST. Finally, 20 sequences were uploaded to NCBI TPA:inferential (Additional file 1: Table S1) as novel chloroplast genomes. Moreover, a search for the species name unveiled that 7 of the 20 sequences are used as ornamental plant, in folk medicine, or as crop plant.

**Table 4.**
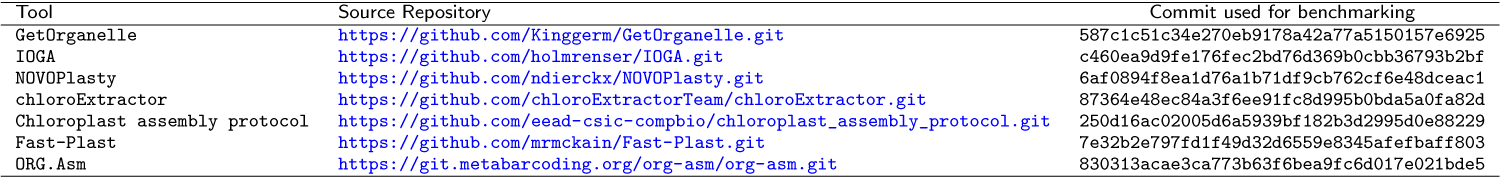
Tools and version information used in our benchmark setup. All tools are wrapped into docker containers and stored on dockerhub [52]. The corresponding tags and SHA256 checksums are reported in Additional file 1: Table S3

**Table 5.** Number of distinct features in novel chloroplast genomes. The distribution (mean, standard deviation (SD), medain, interquartile range (IQR)) of feature types tRNA, rRNA, and coding sequence (CDS) are listed separately.

| Feature | Mean | SD | Median | IQR |
| --- | --- | --- | --- | --- |
| CDS | 79.1 | 3.45 | 80 | 2 |
| rRNA | 4.2 | 0.37 | 4 | 0 |
| tRNA | 26.7 | 0.97 | 27 | 0 |

## Discussion

We compared the overall performance of the different chloroplast assemblers. Depending on the type of downstream applications, the various assessment criteria, need to be weighted differently. For example, ease of installation and use might not be a big concern if the tool is installed once and integrated in an automated pipeline. On the other hand this factor alone might prevent users from being able to use the respective tool in the first place. Similarly, computational requirements or run time might be less relevant, if the goal is to assemble a single chloroplast for further analysis, but are essential if hundreds or thousands of samples will be processed in parallel for a large scale study. Ultimately, both ease of use and computational requirements are irrelevant, if the tool is not able to successfully produce reliable assemblies.

All tools were evaluated under the assumption that they are used in their most basic form (e.g. using default parameters, no pre-processing of the data or post-processing of the result). It is important to note that any tool might perform significantly differently, if distinct parameters are specifically fine-tuned for each data set.

The best performance overall, both on simulated and real data, was achieved by `GetOrganelle`. `Fast-Plast` performed nearly as well on most data. Both tools complement each other, as one tool can achieve successful assemblies of full chloroplasts in cases where the other tool fails. This is highlighted by looking at the de-novo assemblies of chloroplasts, where `GetOrganelle` managed to generate assemblies for 15 different data sets, where no other tool succeeded and `Fast-Plast` was able to assemble 3 plastid genomes that defeated all other tools. `NOVOPlasty` was the only other tool, that could produce an assembly that was not generated with any other assembler. `Fast-Plast`, `NOVOPlasty`, and `ORG.Asm` produced the most variable results, and therefore re-running the tool after a failed attempt might be a valid strategy. `chloroExtractor` yielded only few complete chloroplast assemblies, but requires few resources and is easy to install and use. Thus `chloroExtractor` could be considered as a good option for a quick first try. Both `IOGA` and `Chloroplast assembly protocol` had unsatisfactory performances and failed to return reliable chloroplast assemblies. Nevertheless, multiple alignments of the assembled chloroplast genomes revealed some common challenges for the different tools. Those challenges include fragmented assemblies, invertions of the SSC, or a changed location of the IR (Figure 9).

**Figure 9.**
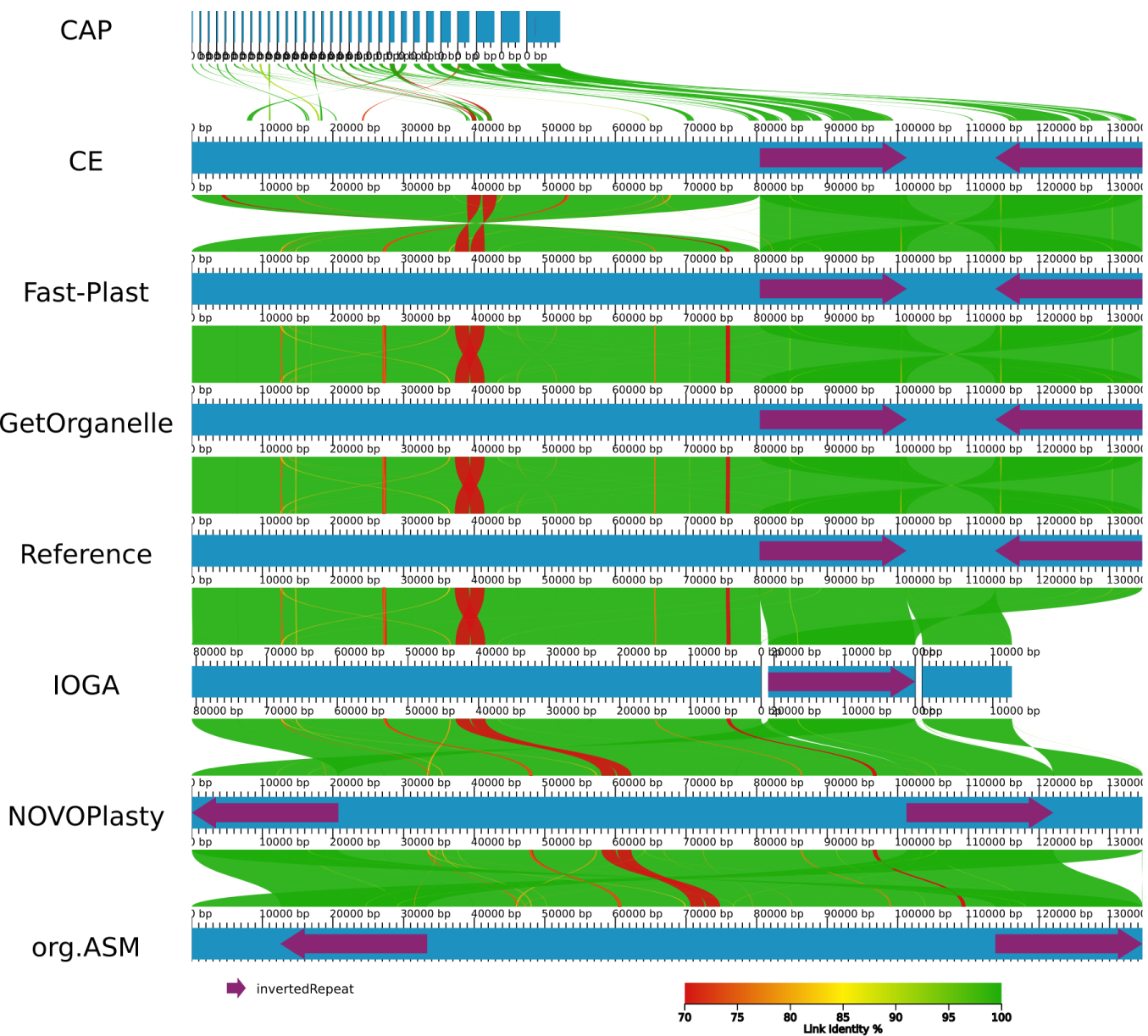
AliTV plot [47] showing alignments of the different assemblers compared to each other and the reference. Each blue bar in the plot corresponds to an assembly of the *Oryza brachyantha* chloroplast (DRR053294). Regions in adjacent assemblies are connected with colored ribbons for similar regions in the alignment with identity coded as color from red to green. The purple arrows on the assemblies depict the IR regions. This plot highlights some common problems with chloroplast assembly. `Chloroplast assembly protocol` only assembled small fragments mostly from the IR region. IOGA returned three separate contigs corresponding to LSC, SSC, and IR. `Fast-Plast` and `GetOrganelle` produced assemblies that were identical to the reference. `chloroExtractor` has the same structure but the LSC is flipped compared to the reference (which is a biologically valid option). Both `NOVOPlasty` and `ORG.Asm` had the SSC flipped compared to the reference and did not start with the LSC but the IR and SSC, respectively (the start point is arbitrary in a circular sequence).

**Figure 10.**
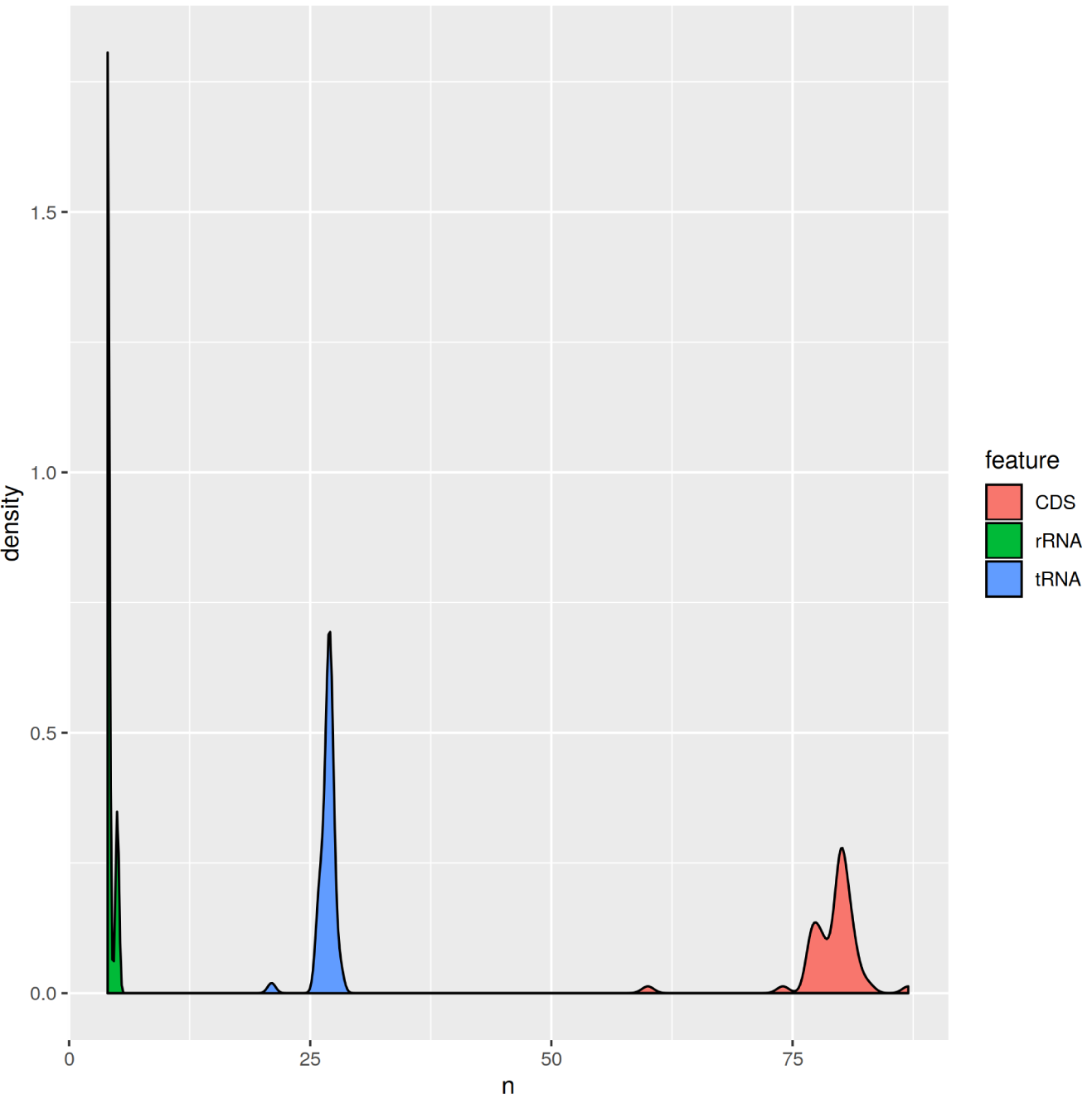
Number of distinct features in novel chloroplast genomes. The distribution of feature types tRNA, rRNA, and coding sequence (CDS) are shown separately.

We observed no phylogenetic pattern in the success rate of the assemblers (Figure 7). This indicates that the tools are generally able to reconstruct chloroplast genomes across the plant kingdom even without available reference genomes.

### Guidelines for the end-user

Given these results, our recommendation is to use `GetOrganelle` as a default option for chloroplast assemblies. If `GetOrganelle` does not produce a use-able assembly, `Fast-Plast` is a valid back-up solution that might be successful. This procedure maximizes the chance of effectively and efficiently recovering the circular chloroplast genome. If both programs fail, it is recommended to try `NOVOPlasty` or manually fine-tune the parameters of the different tools. It is obviously not possible to provide general guidelines, as the exact procedure will differ for different data sets.

For an automated approach, running `GetOrganelle` and `Fast-Plast` in parallel appears to be a good trade-off between success rate and use of resources.

### Ideas for future development

For further experiments, combining different components from different tools might be a promising approach. For example, read scaling from `chloroExtractor` followed by an assembly by `GetOrganelle` and finally structural resolution with `Fast-Plast` could be a promising approach, combing the respective strengths of the different tools.

Moreover, the installation issues need to be mitigated by modern software. Therefore, either containerization (docker, singularity, etc.) or install workflows (e.g. bioconda [50]) should be established by all software packages. Otherwise, the burden of the software installation might result in a low level of uptake by the research community.

A comprehensive documentation, which needs to be up-to-date and maintained, is another important feature of good tools.

All tools should improve their integrated guessing of default parameters, as these are seldom fine-tuned by users, and especially for larger screening approaches.

Finally, as sequencing technology is developing fast (e.g. PacBio or Nanopore), tools need to be updated to be able to handle this new generation of sequencing data and to not become obsolete. The hope would be that with ongoing software development and improved sequencing technologies, the generation of whole chloroplast assemblies from any species will become a routine technique.

## Conclusion

WGS data are also a rich source for chlorplast assemblies. For nearly half of the analyzed data without available chloroplast genome, we could generate complete assemblies using at least one of the tools.

Still, even with simulated (i.e. “perfect”) data, not all tools succeeded in generating complete chloroplast assemblies. Therefore, we determined the strengths and weaknesses of the specific tools and have provided guidelines for users. It might however be necessary to combine different methods or manually explore the parameter space. Ultimately, large scale studies reconstructing hundreds or thousands of chloroplast genomes are now feasible using the currently available tools.

## Methods

### Data availability

Source code for all methods used is available at [51] and archived in Zenodo under [49]. All used assembly tools are hosted on GitHub (Table 4) and are encapsulated in docker containers. That docker containers are published on dockerhub [52] and are named with a leading `benchmark` (Additional file 1: Table S3).

To enable a fair comparison of all tools, we generated simulated sequencing data. Those simulated data sets are stored at Zenodo [53]. All resulting assemblies are available from Zenodo [54]. This study adheres to the guidelines for computational method benchmarking [55].

### Tool Selection

We included tools designed for assembling chloroplasts from whole genome paired end Illumina sequencing data. As a requirement, all tools had to be available as open source software and allow execution via a command line interface. As a graphical user interface (GUI) is not suitable for automated comparisons, tools that only provided a graphical interface were also excluded. The following tools were determined to be within the scope of this study: `ORG.Asm` [27], `chloroExtractor` [33], `Fast-Plast` [56], `IOGA` [57], `NOVOPlasty` [34], `GetOrganelle` [58], and `Chloroplast` `assembly protocol` [59].

Some other related tools for assembling chloroplasts that did not meet our criteria and were therefore outside the scope of this study include: `Organelle PBA` [60]; `sestaton/Chloro` [61]; `Norgal` [62]; `MitoBim` [63].

`Organelle PBA` is designed for PacBio data and does not work with paired Illumina data alone. `sestaton/Chloro` fits our criteria, but is flagged as a work in progress and development and support seem to have ended two years ago. `Norgal` is a tool to extract organellar DNA from whole genome data based on a *k* -mer frequency approach. The final output is a set of contigs of mixed mitochondrial and plastid origin, however. The suggested approach to get a finished chloroplast genome is to run `NOVOPlasty` on the ten longest contigs. We therefore only included `NOVOPlasty` with the default settings and excluded `Norgal`. `MitoBim` is specifically designed for mitochondrial genomes. Even though there is a claim by the author that it can also be used for chloroplasts, there is no further description on how to do so [64].

Additionally, there is a protocol for the `Geneious` [65] software available [66]. However, `Geneious` is closed source and GUI based, which was not in the scope of this study. There is also another publication describing a method for assembling chloroplasts [67]. However, the link to the software is not active anymore.

### Our Setup

We wanted to use a minimum of different parameter settings for all assembly programs to enable a fair comparison. Therefore, we decided to specify that all programs had to work based on two input files, representing the forward (`forward.fq`) and reverse (`reverse.fq`) sequence file of a data set in FASTQ format. Depending on the assembler, output files with different names and locations were generated. Those different files were copied and renamed to ensure that each assembly approach produced the same output file (`output.fa`). Additionally, we set an environment variable for all programs to control the number of allowed threads. All three requirements (defined input file names, defined output file name, thread number control via environment variable) were ensured by a simple wrapper script (`wrapper.sh`). Finally, for a maximum of reproducibility, all programs were bundled into individual docker images based on a central base image which provides all the required software. Those docker images were used for the recording of the consumption of computational resources on a four Intel CPU-E7 8867 v3 system offering 1 TB of RAM. Furthermore, all our docker images have been converted into singularity containers for quantitative measurement on simulated and real data sets. Singularity container were built from docker images for usage on an HPC-environment using Singularity v.2.5.2 [68]. All singularity containers were run on Intel® Xeon® Gold 6140 Processors using a Slurm workload manager version 17.11.8 [69]. Assemblies were run on 4 threads using 10 GiB RAM with a time limit of 48 h.

### Data

#### Simulated data

To avoid complications from sequencing errors and biological variation, we simulated perfect reads based on the *A. thaliana* (TAIR10) chloroplast and core genome assembly [70]. We used a sliding window approach with `seqkit` [71]. The exact commands are documented in `03_representative datasets.md` in [53]. For the final simulated data sets reads are based on the TAIR10 reference genome. Different ratios between the *A. thaliana* core genome in combination with its mitochondrial sequence and the chloroplast sequence were generated (0:1, 1:10, 1:100, and 1:1000). The final data contained 30 × genome coverage and 300 × mitochondrial coverage, except the 0:1 ratio. Additionally, we generated data with different read lengths (150 bp and 250 bp). We further sampled each data set to create another version containing exactly 2 million read pairs.

#### Real data

We selected real data deposited at SRA [26]. We searched all data that matched `(((((((“green plants”[orgn]) AND “wgs”[Strategy]) AND “illumina”[Platform]) AND “biomol dna”[Properties]) AND “paired”[Layout]) AND “random”[Selection]))` `AND “public”[Access]`[72]. For each species with a reference chloroplast in CpBase [73], we selected one data set. In total, this amounted to 369 data sets (Table S2) representing a broad spectrum of the green plants (Figure 7).

#### Novel data

To evaluate the performance for chloroplasts without a reference in CpBase [73], we sampled 105 data sets from the SRA [26] real data set described above (Additional file 1: Table S7). For each entry within that novel data set the number of lineage splits between the source taxon and the related references from CpBase was calculated according to NCBI Taxonomy [74]. The final successful assembly of 49 new chloroplasts was manually inspected and rotated to follow the expected orientation and order of chloroplast genome parts. Due to a lack of a clear definition, we followed the definition of `Fast-Plast` [75].

### Evaluation Criteria

#### Computational Resources

We recorded the mean and the peak CPU usage, the peak memory consumption, and the size of the assembly folder for each program. As input data, we used different data sets comprising 25 000, 250 000 and 2 500 000 read pairs sampled from our simulated reads. We used our docker image setup (Additional file 1: Table S3) to run all assembly programs three times for each parameter setting. The different settings combined different input data and different number of threads to use (1, 2, 4 and 8).

Some programs want to use more CPU threads than specified, therefore, the number of CPUs available was limited using the `--cpu` option of the corresponding `docker run` command. For each assembly setting, we recorded the peak memory consumption, the CPU usage (mean and peak CPU usage) and the size of the folder where the assembly was calculated. The values of CPU and memory usage were obtained from docker. The disk usage was estimated using the GNU tool `du`. We used `GNU parallel` for queuing of the different settings [76].

#### Qualitative

The qualitative evaluation was mainly based on the reviewer guidelines for the Journal of Open Source Software (JOSS) [77]. To create a standard environment, all tools were tested in a fresh default installation of Ubuntu 18.04.2 running in a virtual machine (VirtualBox Version 5.2.18 Ubuntu r123745). We chose this setup instead of the docker container, because it resembles a typical user environment better than the minimal docker installation. The tools were installed according to their installation instructions and the provided tutorial or example usage was executed. During the evaluation, the following questions were asked: (1) Is the tool easy to install? (2) Is there a way to test the installation or a tutorial on how to use the tool? (3) Is there good documentation of the parameter settings? (4) Is the tool maintained (issues answered, implementation of new features)? (5) Is the tool Open Source?

These questions were subjectively answered with Good, Okay or Bad, depending on the quality of the result. For example, a Good installation utilized an automated package or dependency management like `apt`, CRAN, docker, etc. An Okay installation procedure provided a custom script to install everything or at least list all dependencies. A Bad installation procedure failed to list important dependencies or produced errors, that prevented a successful installation without extensive debugging.

After an initial evaluation, we contacted all authors via their GitHub or GitLab issue tracking to communicate potential flaws we found.

#### Quantitative

For each data set and assembler the generated chloroplast genome was compared to the respective reference genome using a pairwise alignment obtained with `minimap2` v2.16 [78]. Based on theses alignments a score was calculated (eq. (1)). The assemblies were scored on a scale from 0 to 100, with 100 being the best and 0 the worst possible score. Four different metrics were incorporated, each contributing a quarter to the total score: completeness; correctness; repeat resolution; continuity. These metrics are similar in concept by those used in the Assemblathon 2 project: coverage; validity; multiplicity; parsimony [79].

The completeness was estimated as the coverage of the assembled chloroplast genome versus the reference genome (*cov*_*ref*_). It represents how many bases of the query genome can be mapped to its respective reference genome. Secondly, we mapped the reference genome against the query. The coverage of the reference genome (*cov*_*qry*_) was used as a measurement of the correctness of the assembly. The repeat resolution was estimated from the size difference of the assembly and the reference genome 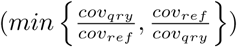, leading to values between 0 and 1. The fourth metric used was the continuity, represented by the number of contigs. A perfect score was achieved if one circular chromosome was assembled, while the score became worse as the number of contigs increased.

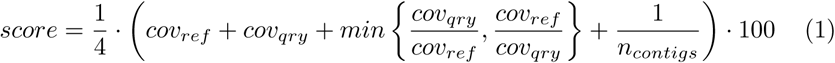

#### Success

For assemblies with reference sequence we defined success as reaching a score of 99 or higher. For the novel chloroplasts our score could not be calculated. The following criteria were instead selected to classify a novel chloroplast as success: single contig of length at least 130 kbp and an IR region of at least 17 kbp. These cutoffs were selected as they produced the highest f-score on the real data set where true assignment (success/failure) was assumed based on the score (success if score 99 or higher).

#### Consistency

To ensure consistency of the obtained results, we rerun and re-evaluated all the assemblies. The resulting assemblies were scored again as described and the scores of the first and the second run were compared to each other This information was important to assess the robustness of the different programs.

## Competing interests

Authors SP, NT, FF, and MJA are developers of chloroExtractor, one of the tools benchmarked in this article. JF, NT, and MJA are affiliated with the for-profit organization AnaLife Data Science.

## Author’s contributions

MJA and FF conceived the project and supervised the findings. SP and FF created the docker images. NT performed the qualitative analysis for all assemblers. MJA prepared the simulated and real data sets. JAF assembled the real data sets. FF ran the performance assemblies on the simulated data sets. All authors developed the score model. JAF and MJA implemented the score model and prepared the figures. All authors discussed the results and contributed to the final manuscript.

## Acknowledgement

We thank Brooke Morriswood for proof-reading and English editing of the manuscript

## Funding

Not applicable.

## Ethics approval

Not applicable.

## Availability of data and materials

The supplemental material is available from Zenodo [80]. The simulated data set is available from Zenodo [53]. All program code is available via Zenodo [49] or from Github [51]. The input data sets can be generated using the raw reads from NCBI SRA (links for each data set in Table S2). The resulting assemblies are available from Zenodo [54].

## Additional Files

Additional file 1 — supplemental data

Supplementary data contain a complete list of all real data sets used in this study. Additionally, a table with more details on the used docker images and the detailed results of the performance measurement are included. The file is available at [80].

## Notes

http://doi.org/10.5281/zenodo.3993699

https://github.com/chloroExtractorTeam/benchmark

http://doi.org/10.5281/zenodo.3836885

### Summary of Updates

Included minor reviewer comments

