## Additional file 1 for "A systematic comparison of chloroplast genome assembly tools"

**Supplemental material**

Jan A Freudenthal, Simon Pfaff, Niklas Terhoeven, Arthur Korte, Markus J Ankenbrand and Frank Förster

Table S1: Sequencing Read Archive and Genbank accessions for novel chloroplast genomes

| Species | SRA-Accession | Assembler | Uses | Genbank-Accession |
|---|---|---|---|---|
| *Antirrhinum majus* subsp. *Striatum* | ERR2744402 | `GetOrganelle` | ornamental plant | - |
| *Antirrhinum majus* var. *pseudomajus* | ERR2744401 | `GetOrganelle` | ornamental plant | - |
| *Arachis magna* | SRR8481359 | `GetOrganelle` | - | - |
| *Boechera stricta* | SRR6821984 | `NOVOPlasty` | - | - |
| *Buddleja globosa* | SRR6331521 | `GetOrganelle` | folk medicine | - |
| *Congea tomentosa* | SRR6940055 | `GetOrganelle` | - | - |
| *Cucurbita argyrosperma* | SRR2531270 | `GetOrganelle` | crop plant | - |
| *Cucurbita ecuadorensis* | SRR2531278 | `GetOrganelle` | - | - |
| *Cucurbita lundelliana* | SRR2531284 | `GetOrganelle` | - | - |
| *Cucurbita okeechobeensis* | SRR2531280 | `GetOrganelle` | - | - |
| *Cucurbita pedatifolia* | SRR2531289 | `GetOrganelle` | - | - |
| *Cuscuta campestris* | ERR1917165 | `GetOrganelle` | - | - |
| *Erysimum baeticum* | SRR7223704 | `GetOrganelle` | - | - |
| *Erysimum nevadense* | SRR7223710 | `GetOrganelle` | - | - |
| *Garrya flavescens* | SRR2401791 | `GetOrganelle` | - | - |
| *Gerbera hybrid cultivar* | SRR2154064 | `GetOrganelle` | - | - |
| *Leucanthemum vulgare* | ERR3089127 | `GetOrganelle` | ornamental plant | - |
| *Petraeovitex bambusetorum* | SRR6940081 | `GetOrganelle` | - | - |
| *Pityopsis graminifolia* var. *Graminifolia* | SRR4457826 | `GetOrganelle` | ornamental plant | - |
| *Pityopsis graminifolia* var. *tenuifolia* | SRR4457835 | `GetOrganelle` | ornamental plant | - |

Table S2: **Summary real data set.** List of real data sets used during our benchmark study. The tables was sorted by species name. Additionally, the table contains the accession for the Sequencing Read Archive (SRA Acc) and the accession of the used reference (Ref Acc). Moreover, the original number of read pairs (read pairs), the overall number of sequenced nucleotides (base pairs), and the mean read length (read len.) is given. Finally, the last column contains the information, if that data set was used for the check of the consistency.

| Taxon | SRA Acc | Ref Acc | read pairs | base pairs (bp) | read len. (bp) | Consistency check |
|---|---|---|---|---|---|---|
| *Abelmoschus esculentus* | SRR5812498 | NC_035234 | 9 028 889 | 2 726 724 478 | 151 | Yes |
| *Abies sibirica* | ERR268415 | NC_035067 | 157 369 451 | 31 473 890 200 | 100 | NO |
| *Actinidia chinensis* | DRR083750 | NC_026690 | 1 974 014 | 574 124 022 | 145 | Yes |
| *Aegilops sharonensis* | ERR359708 | NC_024816 | 179 478 974 | 36 254 752 748 | 101 | NO |
| *Aegilops speltoides* | ERR424849 | NC_022135 | 173 225 639 | 34 991 579 078 | 101 | NO |
| *Aegilops tauschii* | SRR091633 | NC_022133 | 114 178 484 | 23 064 053 768 | 101 | NO |
| *Ajuga reptans* | SRR6940062 | NC_023102 | 16 864 855 | 5 059 456 500 | 150 | NO |
| *Aldrovanda vesiculosa* | SRR7072768 | NC_035416 | 3 510 392 | 1 060 138 384 | 151 | Yes |
| *Allium cepa* | SRR1686960 | NC_024813 | 90 042 453 | 27 099 552 373 | 150 | Yes |
| *Allium sativum* | SRR5602598 | NC_031829 | 1 270 060 | 748 290 736 | 295 | NO |
| *Alloteropsis angusta* | SRR7528995 | NC_027951 | 18 695 483 | 9 347 741 500 | 250 | NO |
| *Alloteropsis cimicina* | SRR7529015 | NC_027952 | 118 496 918 | 59 248 459 000 | 250 | Yes |
| *Alloteropsis paniculata* | SRR4051980 | NC_032078 | 4 048 486 | 796 542 955 | 98 | NO |
| *Alloteropsis semialata* | SRR7528994 | NC_027824 | 26 426 678 | 13 213 339 000 | 250 | Yes |
| *Aloysia citrodora* | SRR5602597 | NC_034695 | 1 423 723 | 844 386 488 | 297 | NO |
| *Althaea officinalis* | SRR5602596 | NC_034701 | 1 409 711 | 833 584 333 | 296 | NO |
| *Amaranthus hypochondriacus* | SRR3399331 | NC_030770 | 6509 | 3 113 188 | 239 | NO |
| *Ammopiptanthus nanus* | SRR6175451 | NC_034743 | 189 485 569 | 56 845 670 700 | 150 | NO |
| *Anacardium occidentale* | SRR5812497 | NC_035235 | 27 997 137 | 7 055 278 524 | 126 | NO |
| *Apostasia odorata* | SRR6037860 | NC_030722 | 48 546 649 | 14 563 994 700 | 150 | Yes |
| *Arabidopsis arenicola* | DRR054584 | NC_030346 | 31 543 276 | 6 371 741 752 | 101 | Yes |
| *Arabidopsis arenosa* | SRR4128972 | NC_029334 | 129 638 900 | 38 891 670 000 | 150 | Yes |

Continued on next page

Table S2 – *Continued from previous page*

| Taxon | SRA Acc | Ref Acc | read pairs | base pairs (bp) | read len. (bp) | Consistency check |
|---|---|---|---|---|---|---|
| *Arabidopsis cebennensis* | SRR2040775 | NC_029335 | 20 358 302 | 4 112 377 004 | 101 | Yes |
| *Arabidopsis croatica* | SRR7637319 | NC_030347 | 14 265 071 | 3 573 463 604 | 125 | NO |
| *Arabidopsis halleri* | SRR8040824 | NC_034366 | 128 303 531 | 38 747 666 362 | 151 | Yes |
| *Arabidopsis lyrata subsp. lyrata* | SRR1705621 | NC_034379 | 16 771 948 | 4 796 683 328 | 143 | NO |
| *Arabidopsis lyrata* | SRR8157563 | NC_034365 | 46 678 588 | 9 429 074 776 | 101 | NO |
| *Arabidopsis neglecta* | SRR2040831 | NC_030348 | 52 410 983 | 10 482 196 600 | 100 | Yes |
| *Arabidopsis pedemontana* | SRR2040802 | NC_029336 | 28 098 774 | 5 675 952 348 | 101 | NO |
| *Arabidopsis petrogena* | SRR2040807 | NC_030349 | 32 189 424 | 6 502 263 648 | 101 | Yes |
| *Arabidopsis suecica* | SRR2084154 | NC_030350 | 158 377 327 | 31 675 465 400 | 100 | NO |
| *Arabidopsis thaliana* | SRR3340908 | NC_000932 | 6 829 274 | 4 097 564 400 | 300 | NO |
| *Arabidopsis umezawana* | SRR2040810 | NC_030351 | 24 335 095 | 4 915 689 190 | 101 | NO |
| *Arabis alpina* | SRR7880736 | NC_023367 | 12 603 925 | 3 126 123 117 | 124 | Yes |
| *Artemisia annua* | SRR5602595 | NC_034683 | 665 200 | 330 541 594 | 248 | Yes |
| *Arundo plinii* | SRR4319202 | NC_034652 | 10 180 635 | 6 099 219 728 | 300 | Yes |
| *Asclepias nivea* | SRR934044 | NC_022431 | 1 637 178 | 311 063 820 | 95 | NO |
| *Asclepias syriaca* | SRR934060 | NC_022432 | 46 704 483 | 9 434 305 566 | 101 | NO |
| *Asparagus officinalis* | DRR056675 | NC_034777 | 83 276 068 | 16 821 765 736 | 101 | Yes |
| *Astragalus mongholicus* | SRR3938254 | NC_029828 | 7 500 181 | 1 514 903 465 | 101 | Yes |
| *Avena sativa* | SRR6056489 | NC_027468 | 115 104 823 | 57 552 411 500 | 250 | NO |
| *Averrhoa carambola* | ERR2799539 | NC_033350 | 1 023 284 | 309 031 768 | 151 | NO |
| *Betula nana* | ERR2026268 | NC_033978 | 45 426 355 | 13 797 661 576 | 152 | NO |
| *Boswellia sacra* | SRR5602594 | NC_029420 | 2 012 947 | 1 199 195 198 | 298 | Yes |
| *Botryococcus braunii* | SRR3721649 | NC_025545 | 137 313 108 | 41 193 932 400 | 150 | NO |
| *Brachypodium distachyon* | SRR4159530 | NC_011032 | 51 410 001 | 15 525 820 302 | 151 | Yes |
| *Brassica juncea* | SRR2057910 | NC_028272 | 54 890 460 | 11 087 872 920 | 101 | NO |
| *Brassica napus* | SRR6856349 | NC_016734 | 34 913 922 | 10 544 004 444 | 151 | NO |
| *Brassica nigra* | SRR2054762 | NC_030450 | 57 198 732 | 11 554 143 864 | 101 | Yes |

Continued on next page

Table S2 – *Continued from previous page*

| Taxon | SRA Acc | Ref Acc | read pairs | base pairs (bp) | read len. (bp) | Consistency check |
|---|---|---|---|---|---|---|
| *Bupleurum latissimum* | SRR6048019 | NC_033346 | 5 702 505 | 3 432 908 010 | 301 | Yes |
| *Cajanus cajan* | SRR073338 | NC_031429 | 132 406 303 | 26 746 073 206 | 101 | Yes |
| *Camelina sativa* | SRR2082664 | NC_029337 | 10 643 512 | 2 149 989 424 | 101 | Yes |
| *Camellia sinensis* | SRR5894861 | NC_020019 | 4 975 058 | 2 985 034 800 | 300 | NO |
| *Cannabis sativa* | SRR7285294 | NC_026562 | 162 968 810 | 42 761 008 291 | 131 | Yes |
| *Capsella bursa-pastoris* | SRR5412136 | NC_009270 | 15 148 015 | 9 391 769 300 | 310 | NO |
| *Capsella grandiflora* | SRR1508428 | NC_028517 | 98 396 864 | 19 679 372 800 | 100 | NO |
| *Capsella rubella* | ERR636124 | NC_027693 | 40 236 079 | 12 151 295 858 | 151 | NO |
| *Capsicum annuum* | SRR1982881 | NC_018552 | 10 804 729 | 1 966 460 678 | 91 | Yes |
| *Carica papaya* | DRR019500 | NC_010323 | 103 259 953 | 22 510 669 754 | 109 | Yes |
| *Carnegiea gigantea* | SRR5036293 | NC_027618 | 12 642 777 | 7 585 666 200 | 300 | NO |
| *Carthamus tinctorius* | SRR2154065 | NC_030783 | 9 648 870 | 1 949 071 740 | 101 | NO |
| *Castanea mollissima* | SRR8731963 | NC_014674 | 118 123 542 | 35 437 062 600 | 150 | Yes |
| *Catharanthus roseus* | SRR5217671 | NC_021423 | 23 799 598 | 7 139 879 400 | 150 | Yes |
| *Cenchrus americanus* | SRR5204424 | NC_024171 | 149 257 429 | 36 289 690 596 | 122 | Yes |
| *Centaurea diffusa* | SRR2729212 | NC_024286 | 69 630 024 | 13 926 004 800 | 100 | Yes |
| *Chenopodium quinoa* | SRR5938314 | NC_034949 | 41 331 617 | 20 748 471 734 | 251 | NO |
| *Chlamydomonas reinhardtii* | SRR4235199 | NC_005353 | 4 795 246 | 1 448 164 292 | 151 | Yes |
| *Chlorella sorokiniana* | SRR4292644 | NC_023835 | 24 148 422 | 4 877 981 244 | 101 | NO |
| *Chlorella variabilis* | SRR7012756 | NC_015359 | 16 059 428 | 4 837 652 469 | 151 | NO |
| *Chromochloris zofingiensis* | SRR5310953 | NC_029672 | 63 205 006 | 12 641 001 200 | 100 | Yes |
| *Chrysobalanus icaco* | SRR1179655 | NC_024061 | 5 544 469 | 1 119 982 738 | 101 | Yes |
| *Cicer arietinum* | SRR5183095 | NC_011163 | 37 577 869 | 6 764 016 420 | 90 | Yes |
| *Cinnamomum verum* | SRR5812493 | NC_035236 | 4 931 460 | 1 489 300 920 | 151 | NO |
| *Citrullus lanatus* | SRR8523014 | NC_032008 | 27 574 511 | 16 544 706 600 | 300 | Yes |
| *Citrus aurantiifolia* | SRR6188466 | NC_024929 | 132 426 000 | 26 485 200 000 | 100 | NO |
| *Citrus limon* | SRR5602593 | NC_034690 | 1 362 792 | 666 398 607 | 244 | NO |

Continued on next page

Table S2 – *Continued from previous page*

| Taxon | SRA Acc | Ref Acc | read pairs | base pairs (bp) | read len. (bp) | Consistency check |
|---|---|---|---|---|---|---|
| *Citrus maxima* | SRR3407098 | NC_034290 | 149 418 786 | 47 814 011 520 | 160 | NO |
| *Citrus reticulata* | SRR3747540 | NC_034671 | 53 225 257 | 15 967 577 100 | 150 | Yes |
| *Citrus sinensis* | SRR3927459 | NC_008334 | 46 781 458 | 14 034 437 400 | 150 | NO |
| *Coffea arabica* | SRR5602572 | NC_008535 | 2 532 608 | 1 494 738 062 | 295 | NO |
| *Coffea canephora* | ERR321701 | NC_030053 | 33 362 762 | 7 206 356 592 | 108 | Yes |
| *Corymbia torelliana* | SRR4063804 | NC_028410 | 66 804 809 | 20 041 442 700 | 150 | Yes |
| *Couepia guianensis* | SRR1179648 | NC_024063 | 7 411 159 | 1 497 054 118 | 101 | Yes |
| *Cucumis melo subsp. melo* | ERR246536 | NC_015983 | 33 207 205 | 10 094 990 320 | 152 | NO |
| *Cucumis sativus* | SRR6490504 | NC_007144 | 35 348 114 | 10 646 772 194 | 151 | Yes |
| *Cunninghamia lanceolata* | ERR2799542 | NC_021437 | 1 062 614 | 320 909 428 | 151 | Yes |
| *Cymbidium ensifolium* | SRR6117755 | NC_028525 | 12 766 736 | 7 685 575 072 | 301 | Yes |
| *Cymbidium kanran* | SRR6117756 | NC_029711 | 5 534 224 | 3 331 602 848 | 301 | NO |
| *Cymbidium lancifolium* | SRR6117757 | NC_029712 | 5 856 776 | 3 525 779 152 | 301 | NO |
| *Cymbidium macrorhizon* | SRR6117754 | NC_029713 | 5 777 841 | 3 478 260 282 | 301 | NO |
| *Cynodon dactylon* | SRR5457035 | NC_034680 | 121 672 | 65 496 192 | 269 | Yes |
| *Cytinus hypocistis* | ERR964904 | NC_031150 | 8 664 415 | 1 750 211 830 | 101 | NO |
| *Dactylis glomerata* | SRR5236602 | NC_027473 | 131 404 213 | 39 421 263 900 | 150 | NO |
| *Daucus carota* | SRR7601255 | NC_008325 | 208 788 541 | 63 054 139 382 | 151 | Yes |
| *Dendrobium aphyllum* | SRR3932123 | NC_035322 | 24 390 720 | 7 317 216 000 | 150 | Yes |
| *Dendrobium chrysanthum* | SRR3932147 | NC_035336 | 12 371 613 | 3 711 483 900 | 150 | Yes |
| *Dendrobium chrysotoxum* | SRR3932141 | NC_028549 | 12 969 805 | 3 890 941 500 | 150 | Yes |
| *Dendrobium crepidatum* | SRR3932124 | NC_035331 | 10 220 355 | 3 066 106 500 | 150 | NO |
| *Dendrobium denneanum* | SRR3932144 | NC_035324 | 11 765 400 | 3 529 620 000 | 150 | Yes |
| *Dendrobium falconeri* | SRR3932139 | NC_035326 | 11 326 112 | 3 397 833 600 | 150 | NO |
| *Dendrobium nobile* | SRR3932122 | NC_029456 | 11 410 037 | 3 423 011 100 | 150 | Yes |
| *Dendrobium parishii* | SRR3932128 | NC_035339 | 14 183 700 | 4 255 110 000 | 150 | NO |
| *Dendrobium pendulum* | SRR3932138 | NC_029705 | 11 357 541 | 3 407 262 300 | 150 | Yes |

Continued on next page

Table S2 – *Continued from previous page*

| Taxon | SRA Acc | Ref Acc | read pairs | base pairs (bp) | read len. (bp) | Consistency check |
|---|---|---|---|---|---|---|
| *Dendrobium primulinum* | SRR3932120 | NC_035321 | 10 864 205 | 3 259 261 500 | 150 | NO |
| *Dendrobium wardianum* | SRR3932135 | NC_035329 | 9 367 500 | 2 810 250 000 | 150 | Yes |
| *Deschampsia antarctica* | SRR1158316 | NC_023533 | 153 346 825 | 30 976 058 650 | 101 | Yes |
| *Digitalis lanata* | SRR5602573 | NC_034688 | 1 257 524 | 730 269 772 | 290 | NO |
| *Dionaea muscipula* | SRR7072324 | NC_035417 | 16 215 933 | 4 897 211 766 | 151 | NO |
| *Dioscorea rotundata* | DRR063110 | NC_024170 | 18 637 125 | 9 295 053 322 | 249 | NO |
| *Dioscorea villosa* | SRR5602590 | NC_034686 | 1 447 023 | 858 275 557 | 297 | Yes |
| *Diospyros lotus* | SRR1560932 | NC_030786 | 8 240 803 | 1 648 160 600 | 100 | Yes |
| *Drosera erythrorhiza* | SRR7072766 | NC_035241 | 5 832 398 | 1 761 384 196 | 151 | NO |
| *Drosera regia* | SRR7072322 | NC_035415 | 1 341 304 | 405 073 808 | 151 | NO |
| *Dunaliella salina* | SRR3923695 | NC_016732 | 164 049 728 | 49 214 918 400 | 150 | Yes |
| *Echinacea angustifolia* | SRR5602579 | NC_034324 | 1 669 371 | 878 253 310 | 263 | Yes |
| *Echinacea atrorubens* | SRR5602578 | NC_034323 | 961 923 | 472 814 810 | 246 | NO |
| *Echinacea laevigata* | SRR5602581 | NC_034322 | 1 099 311 | 545 173 423 | 248 | NO |
| *Echinacea pallida* | SRR5602580 | NC_034321 | 2 039 307 | 832 368 111 | 204 | Yes |
| *Echinacea paradoxa* | SRR5602575 | NC_034320 | 3 101 240 | 1 692 835 034 | 273 | NO |
| *Echinacea purpurea* | SRR5602574 | NC_034327 | 5 197 414 | 2 531 457 606 | 244 | Yes |
| *Echinacea sanguinea* | SRR5602577 | NC_034328 | 4 911 880 | 2 224 965 457 | 226 | Yes |
| *Echinacea speciosa* | SRR5602576 | NC_034325 | 970 715 | 483 247 492 | 249 | NO |
| *Echinacea tennesseensis* | SRR5602587 | NC_034326 | 907 178 | 434 759 190 | 240 | NO |
| *Echinochloa crus-galli* | SRR5920285 | NC_028719 | 203 467 365 | 50 866 841 250 | 125 | NO |
| *Echinochloa oryzicola* | SRR5902661 | NC_024643 | 129 112 810 | 32 278 202 500 | 125 | Yes |
| *Elaeis guineensis* | ERR276848 | NC_017602 | 143 340 222 | 28 954 724 844 | 101 | Yes |
| *Eleutherococcus senticosus* | SRR5602586 | NC_016430 | 424 930 | 211 982 751 | 249 | NO |
| *Epimedium koreanum* | SRR8534306 | NC_029943 | 8 308 432 | 1 262 881 664 | 76 | NO |
| *Epimedium sagittatum* | SRR8275049 | NC_029428 | 4 141 262 | 629 471 824 | 76 | NO |
| *Eragrostis tef* | SRR1463402 | NC_029413 | 85 023 843 | 14 454 053 310 | 85 | NO |

Continued on next page

Table S2 – *Continued from previous page*

| Taxon | SRA Acc | Ref Acc | read pairs | base pairs (bp) | read len. (bp) | Consistency check |
|---|---|---|---|---|---|---|
| *Eriobotrya japonica* | SRR5602604 | NC_034639 | 1 852 832 | 918 799 928 | 248 | Yes |
| *Erodium chrysanthum* | SRR576525 | NC_027065 | 31 570 835 | 6 314 167 000 | 100 | NO |
| *Erodium texanum* | ERR2799526 | NC_014569 | 1 158 815 | 349 962 130 | 151 | Yes |
| *Erythranthe lutea* | SRR5307620 | NC_030212 | 10 000 000 | 2 000 000 000 | 100 | Yes |
| *Eucalyptus grandis* | SRR1392700 | NC_014570 | 327 608 152 | 98 282 445 600 | 150 | Yes |
| *Euphorbia esula* | SRR6355713 | NC_033910 | 65 533 012 | 13 106 602 400 | 100 | NO |
| *Fagopyrum tataricum* | SRR5433722 | NC_027161 | 15 222 050 | 6 981 058 500 | 229 | Yes |
| *Festuca arundinacea* | SRR6885354 | NC_011713 | 6 197 329 | 3 085 916 062 | 249 | NO |
| *Ficus carica* | SRR5678803 | NC_035237 | 26 827 466 | 15 073 339 221 | 281 | NO |
| *Ficus racemosa* | SRR1405699 | NC_028185 | 31 025 867 | 4 715 931 784 | 76 | NO |
| *Foeniculum vulgare* | SRR8690417 | NC_029469 | 3 715 298 | 750 490 196 | 101 | NO |
| *Fragaria chiloensis* | SRR1612828 | NC_019601 | 43 583 617 | 10 460 068 080 | 120 | NO |
| *Fragaria vesca subsp. bracteata* | SRR866156 | NC_018766 | 4 311 362 | 819 158 780 | 95 | Yes |
| *Fragaria virginiana* | SRR5602605 | NC_019602 | 1 199 457 | 708 866 398 | 295 | Yes |
| *Francoa sonchifolia* | SRR576529 | NC_021101 | 32 806 903 | 6 561 380 600 | 100 | NO |
| *Geranium maderense* | ERR2799530 | NC_029999 | 1 031 538 | 311 524 476 | 151 | Yes |
| *Ginkgo biloba* | SRR3089810 | NC_016986 | 10 164 078 | 2 541 019 500 | 125 | NO |
| *Glycine dolichocarpa* | SRR1174380 | NC_021648 | 44 850 853 | 9 059 872 306 | 101 | NO |
| *Glycine max* | SRR6796784 | NC_007942 | 70 339 371 | 21 242 490 042 | 151 | NO |
| *Glycine soja* | SRR7725010 | NC_022868 | 131 827 844 | 39 548 353 200 | 150 | Yes |
| *Glycine syndetika* | SRR1176843 | NC_021650 | 51 528 078 | 10 408 671 756 | 101 | NO |
| *Glycine tomentella* | SRR6823488 | NC_021636 | 265 989 287 | 131 814 519 905 | 248 | NO |
| *Glycyrrhiza glabra* | SRR8690419 | NC_024038 | 4 940 810 | 998 043 620 | 101 | Yes |
| *Glyptostrobus pensilis* | ERR2799543 | NC_031354 | 1 061 851 | 320 679 002 | 151 | NO |
| *Gnetum gnemon* | ERR268420 | NC_026301 | 146 146 748 | 29 229 349 600 | 100 | NO |
| *Gossypium anomalum* | SRR3560153 | NC_023213 | 152 611 035 | 30 827 429 070 | 101 | NO |
| *Gossypium arboreum* | SRR2012734 | NC_016712 | 8 946 554 | 5 385 825 508 | 301 | NO |

Continued on next page

Table S2 – *Continued from previous page*

| Taxon | SRA Acc | Ref Acc | read pairs | base pairs (bp) | read len. (bp) | Consistency check |
|---|---|---|---|---|---|---|
| *Gossypium aridum* | SRR8136263 | NC_033396 | 158 194 517 | 31 638 903 400 | 100 | Yes |
| *Gossypium armourianum* | SRR8136258 | NC_033400 | 155 005 858 | 31 001 171 600 | 100 | NO |
| *Gossypium barbadense* | SRR8627357 | NC_008641 | 159 103 409 | 79 869 911 318 | 251 | NO |
| *Gossypium bickii* | SRR3560187 | NC_023214 | 179 898 006 | 35 979 601 200 | 100 | Yes |
| *Gossypium darwinii* | SRR8545538 | NC_016670 | 440 216 228 | 132 945 300 856 | 151 | NO |
| *Gossypium davidsonii* | SRR6334584 | NC_033395 | 117 719 642 | 35 315 892 600 | 150 | NO |
| *Gossypium gossypioides* | SRR3560148 | NC_017894 | 153 288 210 | 30 657 642 000 | 100 | NO |
| *Gossypium harknessii* | SRR8136257 | NC_033333 | 183 115 691 | 36 431 624 874 | 99 | NO |
| *Gossypium herbaceum* | SRR2012721 | NC_023215 | 12 440 256 | 7 489 034 112 | 301 | NO |
| *Gossypium hirsutum* | SRR3560152 | NC_007944 | 4 487 735 | 2 205 113 705 | 246 | Yes |
| *Gossypium klotzschianum* | SRR8136264 | NC_033394 | 161 773 353 | 32 354 670 600 | 100 | NO |
| *Gossypium longicalyx* | SRR617704 | NC_023216 | 534 258 839 | 107 920 285 478 | 101 | Yes |
| *Gossypium mustelinum* | SRR8727328 | NC_016711 | 158 115 596 | 79 374 029 192 | 251 | NO |
| *Gossypium raimondii* | ERR1449077 | NC_016668 | 21 720 967 | 6 559 732 034 | 151 | NO |
| *Gossypium somalense* | SRR3560160 | NC_018110 | 69 625 198 | 13 925 039 600 | 100 | Yes |
| *Gossypium sturtianum* | SRR3560184 | NC_023218 | 95 466 009 | 19 093 201 800 | 100 | NO |
| *Gossypium thurberi* | SRR8076131 | NC_015204 | 181 003 659 | 36 200 731 800 | 100 | Yes |
| *Gossypium tomentosum* | SRR6334536 | NC_016690 | 138 849 836 | 41 654 950 800 | 150 | NO |
| *Gossypium trilobum* | SRR3560157 | NC_033397 | 52 816 676 | 10 563 335 200 | 100 | NO |
| *Gossypium turneri* | SRR8136254 | NC_026835 | 124 131 681 | 37 239 504 300 | 150 | NO |
| *Haberlea rhodopensis* | SRR4428742 | NC_031852 | 8 365 536 | 1 673 107 200 | 100 | NO |
| *Haplostachys haplostachya* | SRR3170741 | NC_029819 | 251 836 | 76 054 472 | 151 | NO |
| *Helianthus annuus* | SRR825830 | NC_007977 | 57 562 191 | 13 124 179 548 | 114 | Yes |
| *Helianthus argophyllus* | SRR2155086 | NC_030275 | 12 784 471 | 2 582 463 142 | 101 | NO |
| *Helianthus debilis* | SRR5907791 | NC_030173 | 7 323 836 | 1 464 767 200 | 100 | NO |
| *Helianthus divaricatus* | SRR3492253 | NC_023109 | 13 246 221 | 2 649 244 200 | 100 | NO |
| *Helianthus giganteus* | SRR5907716 | NC_023107 | 20 626 828 | 4 125 365 600 | 100 | NO |

Continued on next page

Table S2 – *Continued from previous page*

| Taxon | SRA Acc | Ref Acc | read pairs | base pairs (bp) | read len. (bp) | Consistency check |
|---|---|---|---|---|---|---|
| *Helianthus grosseserratus* | SRR5907715 | NC_023108 | 23 480 167 | 4 696 033 400 | 100 | Yes |
| *Heteropogon triticeus* | SRR5446059 | NC_035025 | 14 302 809 | 4 290 842 700 | 150 | Yes |
| *Hibiscus syriacus* | SRR1265942 | NC_026909 | 25 909 841 | 5 233 787 882 | 101 | NO |
| *Hirtella physophora* | SRR1179646 | NC_024066 | 18 840 660 | 3 768 132 000 | 100 | Yes |
| *Hirtella racemosa* | SRR1179649 | NC_024060 | 8 044 385 | 1 624 965 770 | 101 | Yes |
| *Hordeum vulgare subsp. vulgare* | SRR490932 | NC_008590 | 262 845 212 | 65 711 303 000 | 125 | NO |
| *Humulus lupulus* | SRR8690418 | NC_028032 | 6 541 057 | 1 321 293 514 | 101 | Yes |
| *Hydrastis canadensis* | SRR5602606 | NC_034702 | 1 356 811 | 671 823 274 | 248 | Yes |
| *Hyoscyamus niger* | SRR1508958 | NC_024261 | 21 000 000 | 4 200 000 000 | 100 | Yes |
| *Hypseocharis bilobata* | SRR576531 | NC_023260 | 32 280 852 | 6 456 170 400 | 100 | NO |
| *Illicium anisatum* | SRR5602608 | NC_034703 | 3 195 974 | 961 644 484 | 150 | NO |
| *Illicium floridanum* | SRR5602609 | NC_034685 | 1 929 116 | 1 141 799 529 | 296 | NO |
| *Illicium henryi* | SRR5602610 | NC_034699 | 1 240 196 | 611 286 832 | 246 | NO |
| *Illicium verum* | SRR5602607 | NC_034689 | 2 752 799 | 828 427 309 | 150 | NO |
| *Ipomoea batatas* | SRR7868503 | NC_026703 | 55 433 003 | 27 827 367 506 | 251 | NO |
| *Ipomoea nil* | DRR013917 | NC_031159 | 149 492 354 | 44 847 706 200 | 150 | NO |
| *Ipomoea trifida* | SRR6667669 | NC_034670 | 148 892 802 | 47 645 696 640 | 160 | NO |
| *Jasminum sambac* | SRR5602611 | NC_034694 | 4 487 750 | 2 661 867 968 | 297 | NO |
| *Jasminum tortuosum* | SRR5602601 | NC_034691 | 1 468 745 | 726 257 599 | 247 | NO |
| *Jatropha curcas* | DRR001794 | NC_012224 | 99 881 947 | 15 182 055 944 | 76 | NO |
| *Juglans regia* | SRR2057822 | NC_028617 | 111 991 280 | 33 821 366 560 | 151 | Yes |
| *Juniperus cedrus* | SRR1145775 | NC_028190 | 46 193 335 | 9 238 667 000 | 100 | Yes |
| *Juniperus communis* | ERR268423 | NC_035068 | 472 881 723 | 94 576 344 600 | 100 | NO |
| *Lactuca sativa* | SRR8736654 | NC_007578 | 267 081 494 | 80 124 448 200 | 150 | Yes |
| *Lathyrus sativus* | ERR413118 | NC_014063 | 3 876 258 | 783 004 116 | 101 | NO |
| *Laurus nobilis* | SRR5602602 | NC_034700 | 1 774 932 | 880 729 409 | 248 | Yes |
| *Lavandula angustifolia* | SRR5757713 | NC_029370 | 37 808 338 | 11 418 118 076 | 151 | NO |

Continued on next page

Table S2 – *Continued from previous page*

| Taxon | SRA Acc | Ref Acc | read pairs | base pairs (bp) | read len. (bp) | Consistency check |
|---|---|---|---|---|---|---|
| *Lens culinaris* | ERR413115 | NC_027152 | 6 895 415 | 1 392 873 830 | 101 | Yes |
| *Licania alba* | SRR1179652 | NC_024064 | 6 960 753 | 1 406 072 106 | 101 | Yes |
| *Licania sprucei* | SRR1179651 | NC_024065 | 6 492 846 | 1 311 554 892 | 101 | NO |
| *Lilium tsingtauense* | SRR1265940 | NC_027675 | 9 313 992 | 1 881 426 384 | 101 | NO |
| *Liriodendron tulipifera* | SRR4240953 | NC_008326 | 4 000 000 | 1 208 000 000 | 151 | Yes |
| *Litchi chinensis* | SRR5812499 | NC_035238 | 6 720 070 | 2 029 461 140 | 151 | NO |
| *Lotus japonicus* | SRR8115522 | NC_002694 | 175 495 827 | 52 999 739 754 | 151 | Yes |
| *Ludisia discolor* | SRR3484539 | NC_030540 | 14 073 966 | 8 472 527 532 | 301 | Yes |
| *Macadamia integrifolia* | ERR1397263 | NC_025288 | 28 756 805 | 8 684 555 110 | 151 | Yes |
| *Magnolia biondii* | SRR5602588 | NC_034687 | 1 600 124 | 953 606 691 | 298 | NO |
| *Magnolia denudata* | SRR5602603 | NC_018357 | 1 640 979 | 978 183 192 | 298 | Yes |
| *Magnolia officinalis subsp. biloba* | SRR5602589 | NC_020317 | 1 744 003 | 1 040 030 660 | 298 | NO |
| *Malus prunifolia* | SRR3571175 | NC_031163 | 22 532 714 | 6 804 879 628 | 151 | NO |
| *Mangifera indica* | SRR5812496 | NC_035239 | 9 989 213 | 3 016 742 326 | 151 | Yes |
| *Manihot esculenta* | SRR2847384 | NC_010433 | 121 616 638 | 54 727 487 100 | 225 | Yes |
| *Mankyua chejuensis* | SRR7630500 | NC_017006 | 7 279 904 | 4 382 502 208 | 301 | Yes |
| *Marchantia polymorpha* | SRR396657 | NC_001319 | 123 055 214 | 36 916 564 200 | 150 | NO |
| *Medicago truncatula* | SRR1028837 | NC_003119 | 29 663 436 | 8 958 357 672 | 151 | NO |
| *Melianthus villosus* | SRR576532 | NC_023256 | 31 695 892 | 6 339 178 400 | 100 | Yes |
| *Mentha longifolia* | SRR3204556 | NC_032054 | 67 215 785 | 13 577 588 570 | 101 | Yes |
| *Miscanthus sacchariflorus* | SRR559245 | NC_028720 | 73 073 855 | 22 068 304 210 | 151 | NO |
| *Miscanthus sinensis* | SRR4028761 | NC_028721 | 34 003 586 | 10 269 082 972 | 151 | Yes |
| *Mitragyna speciosa* | SRR5602600 | NC_034698 | 1 327 534 | 658 729 522 | 248 | NO |
| *Monsonia emarginata* | ERR2799532 | NC_029694 | 1 089 971 | 329 171 242 | 151 | Yes |
| *Nelumbo lutea* | SRR5115287 | NC_015605 | 45 729 480 | 11 432 370 000 | 125 | Yes |
| *Nelumbo nucifera* | SRR7159815 | NC_025339 | 30 517 828 | 9 155 348 400 | 150 | NO |
| *Nicotiana otophora* | SRR1171700 | NC_032724 | 67 460 219 | 13 626 964 238 | 101 | NO |

Continued on next page

Table S2 – *Continued from previous page*

| Taxon | SRA Acc | Ref Acc | read pairs | base pairs (bp) | read len. (bp) | Consistency check |
|---|---|---|---|---|---|---|
| *Nicotiana tabacum* | SRR954964 | NC_001879 | 792 406 088 | 160 066 029 776 | 101 | Yes |
| *Nicotiana undulata* | SRR8173251 | NC_016068 | 223 212 812 | 45 088 988 024 | 101 | Yes |
| *Ocimum basilicum* | SRR6940087 | NC_035143 | 32 031 837 | 9 609 551 100 | 150 | NO |
| *Oenothera biennis* | SRR1771524 | NC_010361 | 13 426 811 | 2 658 508 578 | 99 | Yes |
| *Oenothera villaricae* | SRR1771538 | NC_030532 | 3 055 389 | 611 077 800 | 100 | NO |
| *Olea europaea subsp. europaea* | ERR375848 | NC_015401 | 27 590 779 | 16 729 938 945 | 303 | NO |
| *Oryza australiensis* | DRR056778 | NC_024608 | 69 350 090 | 20 805 027 000 | 150 | NO |
| *Oryza barthii* | SRR7341625 | NC_027460 | 55 484 960 | 27 742 480 000 | 250 | Yes |
| *Oryza brachyantha* | DRR053294 | NC_030596 | 10 715 682 | 3 214 704 600 | 150 | Yes |
| *Oryza glaberrima* | SRR7341626 | NC_024175 | 62 571 390 | 31 285 695 000 | 250 | NO |
| *Oryza glumipatula* | DRR057991 | NC_027461 | 23 451 341 | 7 035 402 300 | 150 | Yes |
| *Oryza longiglumis* | DRR056658 | NC_034763 | 50 656 466 | 15 196 939 800 | 150 | Yes |
| *Oryza longistaminata* | DRR058031 | NC_027462 | 17 927 163 | 5 378 148 900 | 150 | Yes |
| *Oryza meridionalis* | DRR058012 | NC_016927 | 19 143 143 | 5 742 942 900 | 150 | NO |
| *Oryza meyeriana* | DRR056670 | NC_034765 | 41 115 373 | 12 334 611 900 | 150 | NO |
| *Oryza minuta* | ERR385916 | NC_030298 | 28 110 596 | 6 746 543 040 | 120 | Yes |
| *Oryza officinalis* | DRR000604 | NC_027463 | 25 866 032 | 5 690 527 040 | 110 | Yes |
| *Oryza punctata* | SRR1264539 | NC_027676 | 8 000 000 | 1 600 000 000 | 100 | NO |
| *Oryza ridleyi* | DRR056663 | NC_034764 | 91 020 902 | 27 306 270 600 | 150 | Yes |
| *Oryza rufipogon* | SRR6220521 | NC_017835 | 26 511 359 | 8 006 430 418 | 151 | NO |
| *Oryza sativa Indica Group* | SRR553489 | NC_027678 | 4 956 305 | 2 488 065 110 | 251 | NO |
| *Oryza sativa Japonica Group* | SRR547959 | NC_001320 | 5 918 441 | 2 971 057 382 | 251 | NO |
| *Oryza sativa* | ERR2696318 | NC_031333 | 6 588 721 | 3 947 278 012 | 300 | Yes |
| *Ostreococcus tauri* | SRR4020477 | NC_008289 | 89 130 001 | 17 826 000 200 | 100 | NO |
| *Ostrya rehderiana* | SRR8302715 | NC_028349 | 33 211 296 | 8 145 137 640 | 123 | NO |
| *Panax ginseng* | SRR5533645 | NC_006290 | 105 595 137 | 37 246 922 641 | 176 | Yes |
| *Panax japonicus* | SRR6512791 | NC_028703 | 71 312 188 | 21 020 452 500 | 147 | NO |

Continued on next page

Table S2 – *Continued from previous page*

| Taxon | SRA Acc | Ref Acc | read pairs | base pairs (bp) | read len. (bp) | Consistency check |
|---|---|---|---|---|---|---|
| *Panax notoginseng* | SRR6512795 | NC_026447 | 7 185 943 | 2 100 564 900 | 146 | NO |
| *Panax quinquefolius* | SRR6512793 | NC_027456 | 20 952 746 | 6 152 880 600 | 147 | NO |
| *Panax stipuleanatus* | SRR6512796 | NC_030598 | 13 610 275 | 3 922 500 000 | 144 | Yes |
| *Panicum capillare* | SRR485871 | NC_030493 | 219 099 842 | 43 819 968 400 | 100 | Yes |
| *Panicum miliaceum* | SRR6650558 | NC_029732 | 159 892 965 | 80 266 268 430 | 251 | Yes |
| *Panicum virgatum* | SRR8440701 | NC_015990 | 92 006 218 | 46 187 121 436 | 251 | NO |
| *Papaver somniferum* | SRR6780757 | NC_029434 | 480 182 430 | 252 410 803 808 | 263 | NO |
| *Parinari campestris* | SRR1179645 | NC_024067 | 5 970 365 | 1 206 013 730 | 101 | NO |
| *Paulownia tomentosa* | SRR6940033 | NC_031436 | 29 457 173 | 8 837 151 900 | 150 | NO |
| *Pelargonium citronellum* | SRR576534 | NC_031194 | 30 750 530 | 6 150 106 000 | 100 | Yes |
| *Pelargonium cotyledonis* | ERR2799534 | NC_028052 | 995 842 | 300 744 284 | 151 | NO |
| *Pelargonium x hortorum* | SRR576536 | NC_008454 | 31 358 909 | 6 271 781 800 | 100 | NO |
| *Perilla frutescens* | SRR6940083 | NC_030756 | 34 758 180 | 10 427 454 000 | 150 | Yes |
| *Phaseolus vulgaris* | SRR5807695 | NC_009259 | 17 399 674 | 5 289 500 896 | 152 | Yes |
| *Phoenix dactylifera* | SRR2518264 | NC_013991 | 89 471 372 | 16 641 675 192 | 93 | NO |
| *Phyllostachys edulis* | SRR8245858 | NC_015817 | 196 487 218 | 58 946 165 400 | 150 | NO |
| *Physcomitrella patens* | ERR1638191 | NC_005087 | 97 932 953 | 29 379 885 900 | 150 | Yes |
| *Picea abies* | ERR1727007 | NC_021456 | 2 556 898 | 644 338 296 | 126 | NO |
| *Picea asperata* | ERR1735493 | NC_032367 | 1 862 931 | 469 458 612 | 126 | Yes |
| *Picea crassifolia* | ERR1735613 | NC_032366 | 1 706 374 | 430 006 248 | 126 | NO |
| *Picea glauca* | SRR869482 | NC_028594 | 19 568 105 | 19 568 105 000 | 500 | Yes |
| *Picea jezoensis* | ERR1795324 | NC_029374 | 2 499 244 | 629 809 488 | 126 | Yes |
| *Picea sitchensis* | SRR5028182 | NC_011152 | 28 752 590 | 14 376 295 000 | 250 | Yes |
| *Pimenta dioica* | SRR5602585 | NC_034684 | 1 821 150 | 1 068 525 583 | 293 | Yes |
| *Pinus lambertiana* | SRR2026990 | NC_011156 | 165 888 035 | 49 959 743 872 | 151 | Yes |
| *Pinus massoniana* | SRR7666034 | NC_021439 | 6 755 123 | 2 026 536 900 | 150 | NO |
| *Pinus sylvestris* | ERR268429 | NC_035069 | 162 706 395 | 32 541 279 000 | 100 | NO |

Continued on next page

Table S2 – *Continued from previous page*

| Taxon | SRA Acc | Ref Acc | read pairs | base pairs (bp) | read len. (bp) | Consistency check |
|---|---|---|---|---|---|---|
| *Pinus taeda* | SRR1049756 | NC_021440 | 4 969 100 | 2 534 241 000 | 255 | NO |
| *Piper auritum* | SRR5602592 | NC_034697 | 1 951 892 | 964 587 352 | 247 | NO |
| *Piper nigrum* | SRR5602591 | NC_034692 | 1 342 936 | 797 054 024 | 297 | NO |
| *Pistacia vera* | SRR4453367 | NC_034998 | 62 397 980 | 18 719 394 000 | 150 | Yes |
| *Pisum sativum* | SRR6702454 | NC_014057 | 144 850 725 | 29 259 846 450 | 101 | NO |
| *Podococcus barteri* | SRR2120220 | NC_027276 | 2453 | 671 984 | 137 | Yes |
| *Populus alba* | SRR3678826 | NC_008235 | 40 869 133 | 8 255 564 866 | 101 | Yes |
| *Populus tremula x Populus alba* | SRR1653109 | NC_028504 | 9 920 342 | 2 003 909 084 | 101 | NO |
| *Populus tremula* | ERR1735633 | NC_027425 | 3 234 148 | 815 005 296 | 126 | Yes |
| *Populus trichocarpa* | SRR5467392 | NC_009143 | 298 329 746 | 149 761 532 492 | 251 | NO |
| *Premna microphylla* | SRR6940036 | NC_026291 | 22 624 524 | 6 787 357 200 | 150 | Yes |
| *Primula veris* | SRR1660449 | NC_031428 | 3 263 725 | 1 599 834 089 | 245 | NO |
| *Prunus dulcis* | SRR5602582 | NC_034696 | 1 285 872 | 631 355 142 | 245 | NO |
| *Prunus kansuensis* | SRR3138168 | NC_023956 | 72 248 792 | 14 449 758 400 | 100 | Yes |
| *Prunus mume* | SRR8240066 | NC_023798 | 45 570 976 | 13 671 292 800 | 150 | Yes |
| *Prunus persica* | SRR5073560 | NC_014697 | 9 978 569 | 4 989 866 485 | 250 | NO |
| *Psidium guajava* | SRR5812495 | NC_033355 | 20 356 683 | 5 129 884 116 | 126 | NO |
| *Punica granatum* | SRR5812494 | NC_035240 | 7 814 299 | 2 359 918 298 | 151 | NO |
| *Pyrus pyrifolia* | SRR5196234 | NC_015996 | 33 563 581 | 9 782 203 050 | 146 | Yes |
| *Raphanus sativus* | DRR014095 | NC_024469 | 513 274 067 | 103 681 361 534 | 101 | NO |
| *Rhazya stricta* | ERR351509 | NC_024292 | 36 920 951 | 6 645 771 180 | 90 | NO |
| *Saccharum officinarum* | SRR7771854 | NC_035224 | 91 863 015 | 46 115 233 530 | 251 | Yes |
| *Saccharum spontaneum* | SRR486146 | NC_034802 | 63 408 401 | 19 149 337 102 | 151 | Yes |
| *Salix purpurea* | SRR3927005 | NC_026722 | 191 948 456 | 57 584 536 800 | 150 | Yes |
| *Salix suchowensis* | SRR8255647 | NC_026462 | 53 495 983 | 16 048 794 900 | 150 | Yes |
| *Salvia miltiorrhiza* | SRR1735356 | NC_020431 | 215 771 972 | 43 585 938 344 | 101 | NO |
| *Sanionia uncinata* | SRR6440975 | NC_025668 | 3 961 386 | 2 379 651 487 | 300 | NO |

Continued on next page

Table S2 – *Continued from previous page*

| Taxon | SRA Acc | Ref Acc | read pairs | base pairs (bp) | read len. (bp) | Consistency check |
|---|---|---|---|---|---|---|
| *Schizachyrium scoparium* | SRR6442327 | NC_035032 | 1 476 472 | 442 941 600 | 150 | Yes |
| *Sciadopitys verticillata* | ERR2799544 | NC_029734 | 1 136 673 | 343 275 246 | 151 | Yes |
| *Scutellaria baicalensis* | SRR6940088 | NC_027262 | 27 930 422 | 8 379 126 600 | 150 | Yes |
| *Scutellaria lateriflora* | SRR5602584 | NC_034693 | 1 699 048 | 843 357 767 | 248 | NO |
| *Sequoia sempervirens* | SRR1951920 | NC_030372 | 2 354 579 | 1 412 747 400 | 300 | NO |
| *Sesamum indicum* | SRR3407090 | NC_016433 | 20 637 541 | 5 159 385 250 | 125 | NO |
| *Setaria italica* | ERR2696316 | NC_022850 | 65 844 392 | 19 753 317 600 | 150 | Yes |
| *Setaria viridis* | SRR4063259 | NC_028075 | 106 716 486 | 33 508 976 604 | 157 | Yes |
| *Silene latifolia* | SRR7040334 | NC_016730 | 284 434 421 | 71 108 605 250 | 125 | NO |
| *Solanum berthaultii* | SRR5349620 | NC_034951 | 30 261 020 | 7 565 255 000 | 125 | NO |
| *Solanum commersonii* | SRR5349609 | NC_028069 | 31 268 502 | 7 817 125 500 | 125 | NO |
| *Solanum habrochaites* | DRR098098 | NC_026879 | 95 529 750 | 17 768 533 500 | 93 | NO |
| *Solanum lycopersicum* | DRR022700 | AC_000188 | 4 192 871 | 2 104 821 242 | 251 | Yes |
| *Solanum melongena* | DRR014074 | NC_030207 | 173 060 640 | 34 958 249 280 | 101 | NO |
| *Solanum peruvianum* | SRR1291239 | NC_026881 | 12 772 339 | 2 449 431 901 | 96 | Yes |
| *Solanum pimpinellifolium* | SRR074949 | NC_026882 | 195 678 316 | 39 527 019 832 | 101 | Yes |
| *Solanum tuberosum* | ERR3148763 | NC_008096 | 806 926 | 243 691 652 | 151 | NO |
| *Sorghum bicolor* | SRR3927127 | NC_008602 | 30 065 772 | 9 079 863 144 | 151 | NO |
| *Sorghum timorense* | SRR424217 | NC_023800 | 200 900 821 | 40 180 164 200 | 100 | Yes |
| *Spirodela polyrhiza* | SRR7548934 | NC_015891 | 87 602 747 | 26 401 985 035 | 151 | Yes |
| *Sporobolus michauxianus* | SRR4434178 | NC_029416 | 183 439 649 | 55 031 894 700 | 150 | NO |
| *Stachys byzantina* | SRR3170744 | NC_029825 | 212 918 | 64 301 236 | 151 | Yes |
| *Stachys chamissonis* | SRR3170745 | NC_029822 | 2 570 411 | 776 264 122 | 151 | Yes |
| *Stachys coccinea* | SRR3170746 | NC_029823 | 292 594 | 88 363 388 | 151 | Yes |
| *Stachys sylvatica* | SRR3170747 | NC_029824 | 592 661 | 178 983 622 | 151 | NO |
| *Taraxacum kok-saghyz* | SRR8185394 | NC_032057 | 21 183 799 | 11 049 748 919 | 261 | NO |
| *Taxodium distichum* | ERR2799546 | NC_034941 | 879 574 | 265 631 348 | 151 | Yes |

Continued on next page

Table S2 – *Continued from previous page*

| Taxon | SRA Acc | Ref Acc | read pairs | base pairs (bp) | read len. (bp) | Consistency check |
|---|---|---|---|---|---|---|
| *Taxus baccata* | ERR268425 | NC_035066 | 172 310 721 | 34 462 144 200 | 100 | Yes |
| *Tectona grandis* | SRR6940065 | NC_020098 | 24 194 338 | 7 258 301 400 | 150 | Yes |
| *Tetradesmus obliquus* | ERR1683652 | NC_008101 | 100 944 122 | 20 390 712 644 | 101 | NO |
| *Themeda triandra* | SRR7529014 | NC_035016 | 91 131 777 | 45 565 888 500 | 250 | Yes |
| *Theobroma cacao* | SRR5602583 | NC_014676 | 470 765 | 277 504 475 | 295 | NO |
| *Thlaspi arvense* | SRR1034659 | NC_034362 | 10 761 690 | 5 402 368 380 | 251 | NO |
| *Trifolium subterraneum* | DRR032042 | NC_011828 | 25 964 880 | 15 630 857 760 | 301 | Yes |
| *Triticum aestivum* | SRR5893651 | NC_002762 | 197 767 716 | 104 816 889 480 | 265 | NO |
| *Triticum monococcum* | SRR384895 | NC_021760 | 157 945 836 | 31 905 058 872 | 101 | NO |
| *Triticum urartu* | SRR4010671 | NC_021762 | 123 772 698 | 63 619 166 772 | 257 | NO |
| *Urochloa ruziziensis* | SRR6816710 | NC_030068 | 154 533 231 | 46 669 035 762 | 151 | NO |
| *Utricularia reniformis* | SRR5349713 | NC_029719 | 2 186 073 | 1 157 291 061 | 265 | NO |
| *Vaccinium macrocarpon* | SRR1276173 | NC_019616 | 30 540 329 | 9 223 179 358 | 151 | NO |
| *Vicia sativa* | ERR413103 | NC_027155 | 10 232 390 | 2 046 478 000 | 100 | NO |
| *Vigna unguiculata* | SRR7125688 | NC_018051 | 30 233 396 | 9 070 018 800 | 150 | Yes |
| *Vitis aestivalis* | SRR5891909 | NC_029454 | 17 094 973 | 5 128 491 900 | 150 | Yes |
| *Vitis amurensis* | SRR5891950 | NC_031383 | 21 559 227 | 6 467 768 100 | 150 | NO |
| *Vitis rotundifolia* | SRR5627788 | NC_023790 | 49 224 878 | 14 767 463 400 | 150 | NO |
| *Vitis vinifera* | SRR7160359 | NC_007957 | 8 416 372 | 8 416 372 000 | 500 | NO |
| *Viviania marifolia* | ERR2799536 | NC_023259 | 1 050 522 | 317 257 644 | 151 | NO |
| *Wisteria floribunda* | SRR1265941 | NC_027677 | 9 382 854 | 1 895 336 508 | 101 | Yes |
| *Wollemia nobilis* | SRR1927951 | NC_027235 | 116 590 | 59 624 823 | 256 | NO |
| *Zea mays* | SRR5826129 | NC_001666 | 172 800 442 | 89 856 229 840 | 260 | Yes |

Table S3: Docker images used in our benchmark setup.

| Tool | Image name and tag | SHA256 Checksum |
| --- | --- | --- |
| chloroExtractor | chloroextracteam/benchmark_chloroextractor:v2.0.0 | 8f49e03424b37e699c5fea9391f79dbb9f2d0dc550ce86c4c700f39deec2dacd |
| Chloroplast assembly protocol | chloroextracteam/benchmark_chloroplast_assembly_protocol:v2.0.1 | 452544e3826748af5b988de0cf402bcea6ff5eb17afdcb71931959d2f6270ca6 |
| Fast-Plast | chloroextracteam/benchmark_fastplast:v2.0.0 | a3d06a610f8340ba49c3ff3b27342534e2a5348b17caa4fd3c3f3d327243a272 |
| GetOrganelle | chloroextracteam/benchmark_getorganelle:v2.0.0 | 2ed3a464a82025a196ea56c649d0b0a3472cef76994bb065d56054d93298d956 |
| IOGA | chloroextracteam/benchmark_ioga:v2.0.0 | 4698a21c343c60290bbf16811165a654ff01e8fddeb41cd75d0771b7f45968c0 |
| NOVOPlasty | chloroextracteam/benchmark_novoplasty:v2.0.0 | 106387bad4e8e5c53fb9d4c3bdd60cdddf05a47b52a38697369eeabb947c547c |
| ORG.Asm | chloroextracteam/benchmark_org-asm:v2.0.0 | ff83677c97b7c4e346191b9e04a5162eeea2a8dfd9e60ff84067d33747868f60 |

Table S4: Performance metrics input size 25K

| | program | threads | Peak CPU usage | Peak memory | Peak disk usage (GB) |
|---|---|---|---|---|---|
| 1 | CAP | 1 | 113.32 | 0.31 | 0.06 |
| 2 | CAP | 2 | 166.60 | 0.26 | 0.05 |
| 3 | CAP | 4 | 305.99 | 0.26 | 0.05 |
| 4 | CAP | 8 | 569.12 | 0.37 | 0.06 |
| 5 | CE | 1 | 111.49 | 3.41 | 0.07 |
| 6 | CE | 2 | 199.12 | 3.41 | 0.07 |
| 7 | CE | 4 | 343.22 | 3.41 | 0.07 |
| 8 | CE | 8 | 509.51 | 3.41 | 0.07 |
| 9 | Fast-Plast | 1 | 118.31 | 0.77 | 0.30 |
| 10 | Fast-Plast | 2 | 213.57 | 0.77 | 0.31 |
| 11 | Fast-Plast | 4 | 379.27 | 0.77 | 0.32 |
| 12 | Fast-Plast | 8 | 657.20 | 0.77 | 0.35 |
| 13 | GetOrganelle | 1 | 111.39 | 0.38 | 0.05 |
| 14 | GetOrganelle | 2 | 170.30 | 0.32 | 0.05 |
| 15 | GetOrganelle | 4 | 242.27 | 0.31 | 0.06 |
| 16 | GetOrganelle | 8 | 346.98 | 0.31 | 0.07 |
| 17 | IOGA | 1 | 118.12 | 18.61 | 6.25 |
| 18 | IOGA | 2 | 224.74 | 17.88 | 6.26 |
| 19 | IOGA | 4 | 394.98 | 16.62 | 5.60 |
| 20 | IOGA | 8 | 733.11 | 21.76 | 11.13 |
| 21 | NOVOPlasty | 1 | 78.38 | 0.04 | 0.00 |
| 22 | NOVOPlasty | 2 | 86.64 | 0.04 | 0.00 |
| 23 | NOVOPlasty | 4 | 86.50 | 0.04 | 0.00 |
| 24 | NOVOPlasty | 8 | 87.32 | 0.04 | 0.00 |
| 25 | org.ASM | 1 | 112.08 | 0.79 | 0.03 |
| 26 | org.ASM | 2 | 115.44 | 0.79 | 0.03 |
| 27 | org.ASM | 4 | 105.09 | 0.79 | 0.03 |
| 28 | org.ASM | 8 | 105.78 | 0.79 | 0.03 |

Table S5: Performance metrics input size 250K

| | program | threads | Peak CPU usage | Peak memory | Peak disk usage (GB) |
|---|---|---|---|---|---|
| 1 | CAP | 1 | 114.38 | 0.90 | 0.65 |
| 2 | CAP | 2 | 204.39 | 0.93 | 0.65 |
| 3 | CAP | 4 | 337.09 | 0.97 | 0.65 |
| 4 | CAP | 8 | 652.75 | 1.08 | 0.65 |
| 5 | CE | 1 | 117.57 | 3.96 | 0.63 |
| 6 | CE | 2 | 230.35 | 3.96 | 0.63 |
| 7 | CE | 4 | 414.25 | 3.96 | 0.63 |
| 8 | CE | 8 | 755.76 | 3.97 | 0.63 |
| 9 | Fast-Plast | 1 | 118.81 | 1.40 | 0.78 |
| 10 | Fast-Plast | 2 | 234.12 | 2.25 | 0.78 |
| 11 | Fast-Plast | 4 | 425.89 | 2.32 | 0.79 |
| 12 | Fast-Plast | 8 | 845.94 | 2.52 | 0.79 |
| 13 | GetOrganelle | 1 | 117.94 | 1.07 | 0.40 |
| 14 | GetOrganelle | 2 | 227.05 | 1.13 | 0.40 |
| 15 | GetOrganelle | 4 | 403.35 | 1.08 | 0.40 |
| 16 | GetOrganelle | 8 | 747.20 | 1.03 | 0.39 |
| 17 | IOGA | 1 | 118.62 | 12.03 | 4.01 |
| 18 | IOGA | 2 | 234.09 | 9.71 | 4.01 |
| 19 | IOGA | 4 | 420.77 | 10.09 | 3.48 |
| 20 | IOGA | 8 | 842.57 | 19.21 | 9.45 |
| 21 | NOVOPlasty | 1 | 111.97 | 0.32 | 0.00 |
| 22 | NOVOPlasty | 2 | 115.07 | 0.32 | 0.00 |
| 23 | NOVOPlasty | 4 | 116.42 | 0.31 | 0.00 |
| 24 | NOVOPlasty | 8 | 115.86 | 0.31 | 0.00 |
| 25 | org.ASM | 1 | 118.10 | 0.93 | 0.06 |
| 26 | org.ASM | 2 | 161.76 | 0.93 | 0.06 |
| 27 | org.ASM | 4 | 163.01 | 0.93 | 0.06 |
| 28 | org.ASM | 8 | 160.74 | 0.93 | 0.06 |

Table S6: Performance metrics input size 2.5M

| | program | threads | Peak CPU usage | Peak memory | Peak disk usage (GB) |
|---|---|---|---|---|---|
| 1 | CAP | 1 | 118.80 | 6.83 | 6.59 |
| 2 | CAP | 2 | 209.22 | 7.58 | 6.59 |
| 3 | CAP | 4 | 357.36 | 7.25 | 6.59 |
| 4 | CAP | 8 | 683.71 | 6.99 | 6.59 |
| 5 | CE | 1 | 118.78 | 5.34 | 2.01 |
| 6 | CE | 2 | 234.82 | 5.34 | 2.01 |
| 7 | CE | 4 | 415.74 | 5.28 | 2.09 |
| 8 | CE | 8 | 844.95 | 5.34 | 2.01 |
| 9 | Fast-Plast | 1 | 119.24 | 6.80 | 5.80 |
| 10 | Fast-Plast | 2 | 237.34 | 6.98 | 5.82 |
| 11 | Fast-Plast | 4 | 443.27 | 8.02 | 5.69 |
| 12 | Fast-Plast | 8 | 872.07 | 8.87 | 5.69 |
| 13 | GetOrganelle | 1 | 118.91 | 5.25 | 3.31 |
| 14 | GetOrganelle | 2 | 235.18 | 5.26 | 3.31 |
| 15 | GetOrganelle | 4 | 420.10 | 5.44 | 3.31 |
| 16 | GetOrganelle | 8 | 845.13 | 6.07 | 3.31 |
| 17 | IOGA | 1 | 119.46 | 18.64 | 8.83 |
| 18 | IOGA | 2 | 237.22 | 21.24 | 10.19 |
| 19 | IOGA | 4 | 425.50 | 18.71 | 10.56 |
| 20 | IOGA | 8 | 857.29 | 25.23 | 16.51 |
| 21 | NOVOPlasty | 1 | 112.67 | 2.69 | 0.00 |
| 22 | NOVOPlasty | 2 | 117.77 | 2.58 | 0.00 |
| 23 | NOVOPlasty | 4 | 117.40 | 2.74 | 0.00 |
| 24 | NOVOPlasty | 8 | 117.28 | 2.74 | 0.00 |
| 25 | org.ASM | 1 | 118.73 | 2.63 | 0.37 |
| 26 | org.ASM | 2 | 184.53 | 2.63 | 0.37 |
| 27 | org.ASM | 4 | 177.03 | 2.63 | 0.37 |
| 28 | org.ASM | 8 | 176.25 | 2.63 | 0.37 |

Table S7: **Novel data set.**

| SRA Accession | Depth | TaxID | Rank | Related References with Depth | Status |
|---|---|---|---|---|---|
| DRR053713 | 3 | 71243 | order | NC_033350(3) | success |
| DRR057122 | 3 | 3650 | family | NC_007144(3) NC_023544(3) NC_029484(3) NC_031834(3) NC_032008(3) NC_033899(3) NC_015983(4) | success |
| DRR106816 | 5 | 2231393 | no rank | NC_009259(5) NC_013843(5) NC_016708(5) NC_018051(5) NC_021091(5) NC_023090(5) NC_025909(5) NC_027677(5) NC_029406(5) NC_031429(5) NC_034742(5) NC_034743(5) NC_034774(5) NC_002694(6) NC_003119(6) NC_007942(6) NC_011163(6) NC_011828(6) NC_014057(6) NC_014063(6) NC_021636(6) NC_021645(6) NC_021646(6) NC_021647(6) NC_021648(6) NC_021649(6) NC_021650(6) NC_022868(6) NC_024034(6) NC_024035(6) NC_024036(6) NC_024038(6) NC_024166(6) NC_025743(6) NC_025744(6) NC_025745(6) NC_027073(6) NC_027074(6) NC_027075(6) NC_027076(6) NC_027077(6) NC_027078(6) NC_027079(6) NC_027080(6) NC_027148(6) NC_027149(6) NC_027150(6) NC_027151(6) NC_027152(6) NC_027153(6) NC_027154(6) NC_027155(6) NC_029828(6) NC_030329(6) NC_032066(6) NC_032691(6) NC_034229(6) NC_035228(6) NC_035229(6) NC_028171(7) | success |
| ERR1917165 | 5 | 4118 | family | NC_009808(3) NC_026703(3) NC_031159(3) NC_034670(3) | success |

Continued on next page

Table S7 – *Continued from previous page*

| SRA Accession | Depth | TaxID | Rank | Related References with Depth | Status |
|---|---|---|---|---|---|
| ERR2003066 | 3 | 721789 | tribe | NC_018767(3) NC_019601(3) NC_019602(3) NC_034347(3) NC_015206(4) NC_018766(4) | success |
| ERR2744401 | 5 | 156152 | family | NC_028519(3) NC_028520(3) NC_031345(3) NC_034688(3) NC_031153(4) NC_031344(4) | success |
| ERR2744402 | 4 | 156152 | family | NC_028519(3) NC_028520(3) NC_031345(3) NC_034688(3) NC_031153(4) NC_031344(4) | success |
| ERR3089127 | 4 | 102810 | tribe | NC_020092(3) NC_020320(3) NC_020607(3) NC_025910(3) NC_030785(3) NC_031399(3) NC_031400(3) NC_034683(3) NC_034851(3) | success |

Continued on next page

Table S7 – *Continued from previous page*

| SRA Accession | Depth | TaxID | Rank | Related References with Depth | | | Status |
|---|---|---|---|---|---|---|---|
| SRR2154064 | 4 | 4210 | family | NC_027835(4) | NC_034648(4) | NC_007578(5) | success |
| | | | | NC_007977(5) | NC_010601(5) | NC_013553(5) | |
| | | | | NC_015543(5) | NC_015621(5) | NC_020092(5) | |
| | | | | NC_020320(5) | NC_020607(5) | NC_023107(5) | |
| | | | | NC_023108(5) | NC_023109(5) | NC_023110(5) | |
| | | | | NC_023111(5) | NC_023112(5) | NC_023113(5) | |
| | | | | NC_023114(5) | NC_023833(5) | NC_024286(5) | |
| | | | | NC_025910(5) | NC_027113(5) | NC_027434(5) | |
| | | | | NC_028005(5) | NC_028006(5) | NC_028027(5) | |
| | | | | NC_029465(5) | NC_030173(5) | NC_030275(5) | |
| | | | | NC_030772(5) | NC_030773(5) | NC_030783(5) | |
| | | | | NC_030785(5) | NC_031395(5) | NC_031396(5) | |
| | | | | NC_031399(5) | NC_031400(5) | NC_031815(5) | |
| | | | | NC_031816(5) | NC_031833(5) | NC_031853(5) | |
| | | | | NC_031898(5) | NC_032056(5) | NC_032057(5) | |
| | | | | NC_034320(5) | NC_034321(5) | NC_034322(5) | |
| | | | | NC_034323(5) | NC_034324(5) | NC_034325(5) | |
| | | | | NC_034326(5) | NC_034327(5) | NC_034328(5) | |
| | | | | NC_034683(5) | NC_034810(5) | NC_034811(5) | |
| | | | | NC_034812(5) | NC_034813(5) | NC_034814(5) | |
| | | | | NC_034815(5) | NC_034816(5) | NC_034817(5) | |
| | | | | NC_034818(5) | NC_034819(5) | NC_034820(5) | |
| | | | | NC_034821(5) | NC_034822(5) | NC_034823(5) | |
| | | | | NC_034824(5) | NC_034825(5) | NC_034827(5) | |
| | | | | NC_034828(5) | NC_034829(5) | NC_034830(5) | |
| | | | | NC_034831(5) | NC_034832(5) | NC_034847(5) | |
| | | | | NC_034848(5) | NC_034849(5) | NC_034850(5) | |
| | | | | NC_034851(5) | NC_034852(5) | NC_034853(5) | |
| | | | | NC_034854(5) | NC_034856(5) | NC_034857(5) | |
| | | | | NC_034858(5) | NC_034859(5) | NC_034860(5) | |
| | | | | NC_034861(5) | NC_034862(5) | NC_034864(5) | |
| | | | | NC_034865(5) | NC_034866(5) | NC_034867(5) | |
| | | | | NC_034868(5) | NC_034869(5) | NC_034870(5) | |
| | | | | NC_034871(5) | NC_034872(5) | NC_034873(5) | |
| | | | | NC_034874(5) | NC_034875(5) | NC_034876(5) | |
| | | | | NC_034877(5) | NC_034878(5) | NC_034879(5) | |
| | | | | NC_034880(5) | NC_034881(5) | NC_034882(5) | |
| | | | | NC_034883(5) | NC_034884(5) | NC_034885(5) | |

Table S7 – *Continued from previous page*

| SRA Accession | Depth | TaxID | Rank | Related References with Depth | Status |
|---|---|---|---|---|---|
| SRR2176133 | 3 | 40550 | order | NC_026786(3) NC_027659(3) | success |

Continued on next page

Table S7 – *Continued from previous page*

| SRA Accession | Depth | TaxID | Rank | Related References with Depth | | | Status |
|---|---|---|---|---|---|---|---|
| SRR2401791 | 4 | 91888 | no rank | NC_016433(4) | NC_021449(4) | NC_023463(4) | success |
| | | | | NC_025652(4) | NC_025653(4) | NC_029719(4) | |
| | | | | NC_030212(4) | NC_031435(4) | NC_031436(4) | |
| | | | | NC_008407(5) | NC_009808(5) | NC_013707(5) | |
| | | | | NC_020098(5) | NC_021111(5) | NC_022859(5) | |
| | | | | NC_023102(5) | NC_023115(5) | NC_023131(5) | |
| | | | | NC_023132(5) | NC_023464(5) | NC_023465(5) | |
| | | | | NC_024845(5) | NC_025641(5) | NC_025642(5) | |
| | | | | NC_025651(5) | NC_025787(5) | NC_026202(5) | |
| | | | | NC_026291(5) | NC_026703(5) | NC_027262(5) | |
| | | | | NC_027838(5) | NC_027955(5) | NC_028519(5) | |
| | | | | NC_028520(5) | NC_028533(5) | NC_029700(5) | |
| | | | | NC_031159(5) | NC_031345(5) | NC_031437(5) | |
| | | | | NC_031441(5) | NC_031442(5) | NC_031443(5) | |
| | | | | NC_031444(5) | NC_031445(5) | NC_033534(5) | |
| | | | | NC_034308(5) | NC_034309(5) | NC_034310(5) | |
| | | | | NC_034311(5) | NC_034312(5) | NC_034670(5) | |
| | | | | NC_034688(5) | NC_034691(5) | NC_034693(5) | |
| | | | | NC_034694(5) | NC_034695(5) | NC_035000(5) | |
| | | | | NC_001879(6) | NC_004561(6) | NC_007500(6) | |
| | | | | NC_007602(6) | NC_007943(6) | NC_008096(6) | |
| | | | | NC_015401(6) | NC_015604(6) | NC_015608(6) | |
| | | | | NC_015623(6) | NC_016068(6) | NC_018117(6) | |
| | | | | NC_018552(6) | NC_022451(6) | NC_024261(6) | |
| | | | | NC_024292(6) | NC_025657(6) | NC_026551(6) | |
| | | | | NC_026563(6) | NC_026567(6) | NC_026570(6) | |
| | | | | NC_026574(6) | NC_026694(6) | NC_026726(6) | |
| | | | | NC_026906(6) | NC_027099(6) | NC_027177(6) | |
| | | | | NC_027441(6) | NC_027442(6) | NC_028007(6) | |
| | | | | NC_028009(6) | NC_028069(6) | NC_028070(6) | |
| | | | | NC_028614(6) | NC_029370(6) | NC_029746(6) | |
| | | | | NC_029817(6) | NC_029818(6) | NC_029819(6) | |
| | | | | NC_029820(6) | NC_029821(6) | NC_029822(6) | |
| | | | | NC_029823(6) | NC_029824(6) | NC_029825(6) | |
| | | | | NC_029833(6) | NC_030044(6) | NC_030056(6) | |
| | | | | NC_030167(6) | NC_030168(6) | NC_030171(6) | |
| | | | | NC_030177(6) | NC_030178(6) | NC_030185(6) | |
| | | | | NC_030207(6) | NC_030282(6) | NC_030319(6) | |

Table S7 – *Continued from previous page*

| SRA Accession | Depth | TaxID | Rank | Related References with Depth | Status |
|---|---|---|---|---|---|
| SRR2531270 | 3 | 3650 | family | NC_007144(3) NC_023544(3) NC_029484(3) NC_031834(3) NC_032008(3) NC_033899(3) NC_015983(4) | success |
| SRR2531276 | 4 | 3650 | family | NC_007144(3) NC_023544(3) NC_029484(3) NC_031834(3) NC_032008(3) NC_033899(3) NC_015983(4) | success |
| SRR2531278 | 3 | 3650 | family | NC_007144(3) NC_023544(3) NC_029484(3) NC_031834(3) NC_032008(3) NC_033899(3) NC_015983(4) | success |
| SRR2531280 | 3 | 3650 | family | NC_007144(3) NC_023544(3) NC_029484(3) NC_031834(3) NC_032008(3) NC_033899(3) NC_015983(4) | success |
| SRR2531284 | 3 | 3650 | family | NC_007144(3) NC_023544(3) NC_029484(3) NC_031834(3) NC_032008(3) NC_033899(3) NC_015983(4) | success |
| SRR2531289 | 3 | 3650 | family | NC_007144(3) NC_023544(3) NC_029484(3) NC_031834(3) NC_032008(3) NC_033899(3) NC_015983(4) | success |

Continued on next page

Table S7 – *Continued from previous page*

| SRA Accession | Depth | TaxID | Rank | Related References with Depth | Status |
|---|---|---|---|---|---|
| SRR4124062 | 7 | 2231393 | no rank | NC_009259(5) NC_013843(5) NC_016708(5) NC_018051(5) NC_021091(5) NC_023090(5) NC_025909(5) NC_027677(5) NC_029406(5) NC_031429(5) NC_034742(5) NC_034743(5) NC_034774(5) NC_002694(6) NC_003119(6) NC_007942(6) NC_011163(6) NC_011828(6) NC_014057(6) NC_014063(6) NC_021636(6) NC_021645(6) NC_021646(6) NC_021647(6) NC_021648(6) NC_021649(6) NC_021650(6) NC_022868(6) NC_024034(6) NC_024035(6) NC_024036(6) NC_024038(6) NC_024166(6) NC_025743(6) NC_025744(6) NC_025745(6) NC_027073(6) NC_027074(6) NC_027075(6) NC_027076(6) NC_027077(6) NC_027078(6) NC_027079(6) NC_027080(6) NC_027148(6) NC_027149(6) NC_027150(6) NC_027151(6) NC_027152(6) NC_027153(6) NC_027154(6) NC_027155(6) NC_029828(6) NC_030329(6) NC_032066(6) NC_032691(6) NC_034229(6) NC_035228(6) NC_035229(6) NC_028171(7) | success |

Continued on next page

Table S7 – *Continued from previous page*

| SRA Accession | Depth | TaxID | Rank | Related References with Depth | Status |
|---|---|---|---|---|---|
| SRR4124063 | 6 | 2231393 | no rank | NC_009259(5) NC_013843(5) NC_016708(5)<br>NC_018051(5) NC_021091(5) NC_023090(5)<br>NC_025909(5) NC_027677(5) NC_029406(5)<br>NC_031429(5) NC_034742(5) NC_034743(5)<br>NC_034774(5) NC_002694(6) NC_003119(6)<br>NC_007942(6) NC_011163(6) NC_011828(6)<br>NC_014057(6) NC_014063(6) NC_021636(6)<br>NC_021645(6) NC_021646(6) NC_021647(6)<br>NC_021648(6) NC_021649(6) NC_021650(6)<br>NC_022868(6) NC_024034(6) NC_024035(6)<br>NC_024036(6) NC_024038(6) NC_024166(6)<br>NC_025743(6) NC_025744(6) NC_025745(6)<br>NC_027073(6) NC_027074(6) NC_027075(6)<br>NC_027076(6) NC_027077(6) NC_027078(6)<br>NC_027079(6) NC_027080(6) NC_027148(6)<br>NC_027149(6) NC_027150(6) NC_027151(6)<br>NC_027152(6) NC_027153(6) NC_027154(6)<br>NC_027155(6) NC_029828(6) NC_030329(6)<br>NC_032066(6) NC_032691(6) NC_034229(6)<br>NC_035228(6) NC_035229(6) NC_028171(7) | success |

Continued on next page

Table S7 – *Continued from previous page*

| SRA Accession | Depth | TaxID | Rank | Related References with Depth | Status |
|---|---|---|---|---|---|
| SRR4124078 | 6 | 2231393 | no rank | NC_009259(5) NC_013843(5) NC_016708(5) NC_018051(5) NC_021091(5) NC_023090(5) NC_025909(5) NC_027677(5) NC_029406(5) NC_031429(5) NC_034742(5) NC_034743(5) NC_034774(5) NC_002694(6) NC_003119(6) NC_007942(6) NC_011163(6) NC_011828(6) NC_014057(6) NC_014063(6) NC_021636(6) NC_021645(6) NC_021646(6) NC_021647(6) NC_021648(6) NC_021649(6) NC_021650(6) NC_022868(6) NC_024034(6) NC_024035(6) NC_024036(6) NC_024038(6) NC_024166(6) NC_025743(6) NC_025744(6) NC_025745(6) NC_027073(6) NC_027074(6) NC_027075(6) NC_027076(6) NC_027077(6) NC_027078(6) NC_027079(6) NC_027080(6) NC_027148(6) NC_027149(6) NC_027150(6) NC_027151(6) NC_027152(6) NC_027153(6) NC_027154(6) NC_027155(6) NC_029828(6) NC_030329(6) NC_032066(6) NC_032691(6) NC_034229(6) NC_035228(6) NC_035229(6) NC_028171(7) | success |

Continued on next page

Table S7 – *Continued from previous page*

| SRA Accession | Depth | TaxID | Rank | Related References with Depth | Status |
| --- | --- | --- | --- | --- | --- |
| SRR4124079 | 7 | 2231393 | no rank | NC_009259(5) NC_013843(5) NC_016708(5)<br>NC_018051(5) NC_021091(5) NC_023090(5)<br>NC_025909(5) NC_027677(5) NC_029406(5)<br>NC_031429(5) NC_034742(5) NC_034743(5)<br>NC_034774(5) NC_002694(6) NC_003119(6)<br>NC_007942(6) NC_011163(6) NC_011828(6)<br>NC_014057(6) NC_014063(6) NC_021636(6)<br>NC_021645(6) NC_021646(6) NC_021647(6)<br>NC_021648(6) NC_021649(6) NC_021650(6)<br>NC_022868(6) NC_024034(6) NC_024035(6)<br>NC_024036(6) NC_024038(6) NC_024166(6)<br>NC_025743(6) NC_025744(6) NC_025745(6)<br>NC_027073(6) NC_027074(6) NC_027075(6)<br>NC_027076(6) NC_027077(6) NC_027078(6)<br>NC_027079(6) NC_027080(6) NC_027148(6)<br>NC_027149(6) NC_027150(6) NC_027151(6)<br>NC_027152(6) NC_027153(6) NC_027154(6)<br>NC_027155(6) NC_029828(6) NC_030329(6)<br>NC_032066(6) NC_032691(6) NC_034229(6)<br>NC_035228(6) NC_035229(6) NC_028171(7) | success |

Continued on next page

Table S7 – *Continued from previous page*

| SRA Accession | Depth | TaxID | Rank | Related References with Depth | Status |
|---|---|---|---|---|---|
| SRR4457826 | 4 | 102809 | tribe | NC_027434(3) NC_034810(3) NC_034811(3) NC_034812(3) NC_034813(3) NC_034814(3) NC_034815(3) NC_034816(3) NC_034817(3) NC_034818(3) NC_034819(3) NC_034820(3) NC_034821(3) NC_034822(3) NC_034823(3) NC_034824(3) NC_034825(3) NC_034827(3) NC_034828(3) NC_034829(3) NC_034830(3) NC_034831(3) NC_034832(3) NC_034847(3) NC_034848(3) NC_034849(3) NC_034850(3) NC_034852(3) NC_034853(3) NC_034854(3) NC_034856(3) NC_034857(3) NC_034858(3) NC_034859(3) NC_034860(3) NC_034861(3) NC_034862(3) NC_034864(3) NC_034865(3) NC_034866(3) NC_034867(3) NC_034868(3) NC_034869(3) NC_034870(3) NC_034871(3) NC_034872(3) NC_034873(3) NC_034874(3) NC_034875(3) NC_034876(3) NC_034877(3) NC_034878(3) NC_034879(3) NC_034880(3) NC_034881(3) NC_034882(3) NC_034883(3) NC_034884(3) NC_034885(3) NC_034886(3) NC_034887(3) NC_034888(3) NC_034889(3) NC_034891(3) NC_034892(3) NC_034893(3) NC_034894(3) NC_034895(3) NC_034896(3) NC_034897(3) NC_034898(3) NC_034899(3) NC_034900(3) NC_034901(3) NC_034902(3) NC_034903(3) NC_034996(3) | success |

Continued on next page

Table S7 – *Continued from previous page*

| SRA Accession | Depth | TaxID | Rank | Related References with Depth | Status |
|---|---|---|---|---|---|
| SRR4457835 | 4 | 102809 | tribe | NC_027434(3) NC_034810(3) NC_034811(3) NC_034812(3) NC_034813(3) NC_034814(3) NC_034815(3) NC_034816(3) NC_034817(3) NC_034818(3) NC_034819(3) NC_034820(3) NC_034821(3) NC_034822(3) NC_034823(3) NC_034824(3) NC_034825(3) NC_034827(3) NC_034828(3) NC_034829(3) NC_034830(3) NC_034831(3) NC_034832(3) NC_034847(3) NC_034848(3) NC_034849(3) NC_034850(3) NC_034852(3) NC_034853(3) NC_034854(3) NC_034856(3) NC_034857(3) NC_034858(3) NC_034859(3) NC_034860(3) NC_034861(3) NC_034862(3) NC_034864(3) NC_034865(3) NC_034866(3) NC_034867(3) NC_034868(3) NC_034869(3) NC_034870(3) NC_034871(3) NC_034872(3) NC_034873(3) NC_034874(3) NC_034875(3) NC_034876(3) NC_034877(3) NC_034878(3) NC_034879(3) NC_034880(3) NC_034881(3) NC_034882(3) NC_034883(3) NC_034884(3) NC_034885(3) NC_034886(3) NC_034887(3) NC_034888(3) NC_034889(3) NC_034891(3) NC_034892(3) NC_034893(3) NC_034894(3) NC_034895(3) NC_034896(3) NC_034897(3) NC_034898(3) NC_034899(3) NC_034900(3) NC_034901(3) NC_034902(3) NC_034903(3) NC_034996(3) | success |

Continued on next page

Table S7 – *Continued from previous page*

| SRA Accession | Depth | TaxID | Rank | Related References with Depth | Status |
|---|---|---|---|---|---|
| SRR5195283 | 3 | 4055 | order | NC_024292(5) NC_025657(5) NC_027441(5) NC_027442(5) NC_028009(5) NC_028614(5) NC_030319(5) NC_031155(5) NC_033354(5) NC_034698(5) NC_021423(6) NC_022431(6) NC_022432(6) NC_025655(6) NC_025656(6) NC_025658(6) NC_029459(6) NC_029460(6) NC_008535(7) NC_030053(7) | success |
| SRR5409224 | 5 | 2231393 | no rank | NC_009259(5) NC_013843(5) NC_016708(5) NC_018051(5) NC_021091(5) NC_023090(5) NC_025909(5) NC_027677(5) NC_029406(5) NC_031429(5) NC_034742(5) NC_034743(5) NC_034774(5) NC_002694(6) NC_003119(6) NC_007942(6) NC_011163(6) NC_011828(6) NC_014057(6) NC_014063(6) NC_021636(6) NC_021645(6) NC_021646(6) NC_021647(6) NC_021648(6) NC_021649(6) NC_021650(6) NC_022868(6) NC_024034(6) NC_024035(6) NC_024036(6) NC_024038(6) NC_024166(6) NC_025743(6) NC_025744(6) NC_025745(6) NC_027073(6) NC_027074(6) NC_027075(6) NC_027076(6) NC_027077(6) NC_027078(6) NC_027079(6) NC_027080(6) NC_027148(6) NC_027149(6) NC_027150(6) NC_027151(6) NC_027152(6) NC_027153(6) NC_027154(6) NC_027155(6) NC_029828(6) NC_030329(6) NC_032066(6) NC_032691(6) NC_034229(6) NC_035228(6) NC_035229(6) NC_028171(7) | success |

Continued on next page

Table S7 – *Continued from previous page*

| SRA Accession | Depth | TaxID | Rank | Related References with Depth | Status |
|---|---|---|---|---|---|
| SRR5409225 | 4 | 2231393 | no rank | NC_009259(5) NC_013843(5) NC_016708(5) NC_018051(5) NC_021091(5) NC_023090(5) NC_025909(5) NC_027677(5) NC_029406(5) NC_031429(5) NC_034742(5) NC_034743(5) NC_034774(5) NC_002694(6) NC_003119(6) NC_007942(6) NC_011163(6) NC_011828(6) NC_014057(6) NC_014063(6) NC_021636(6) NC_021645(6) NC_021646(6) NC_021647(6) NC_021648(6) NC_021649(6) NC_021650(6) NC_022868(6) NC_024034(6) NC_024035(6) NC_024036(6) NC_024038(6) NC_024166(6) NC_025743(6) NC_025744(6) NC_025745(6) NC_027073(6) NC_027074(6) NC_027075(6) NC_027076(6) NC_027077(6) NC_027078(6) NC_027079(6) NC_027080(6) NC_027148(6) NC_027149(6) NC_027150(6) NC_027151(6) NC_027152(6) NC_027153(6) NC_027154(6) NC_027155(6) NC_029828(6) NC_030329(6) NC_032066(6) NC_032691(6) NC_034229(6) NC_035228(6) NC_035229(6) NC_028171(7) | success |
| SRR5581778 | 4 | 3650 | family | NC_007144(3) NC_023544(3) NC_029484(3) NC_031834(3) NC_032008(3) NC_033899(3) NC_015983(4) | success |
| SRR5586278 | 5 | 1521262 | subclass | NC_012818(4) NC_022136(4) NC_022137(4) NC_024157(4) NC_024158(4) NC_004766(6) NC_014348(6) NC_014592(6) NC_028542(6) NC_028543(6) NC_028705(6) | success |

Continued on next page

Table S7 – *Continued from previous page*

| SRA Accession | Depth | TaxID | Rank | Related References with Depth | | | Status |
|---|---|---|---|---|---|---|---|
| SRR5880481 | 3 | 41945 | order | NC_020019(3) | NC_021121(3) | NC_022264(3) | success |
| | | | | NC_022459(3) | NC_022460(3) | NC_022461(3) | |
| | | | | NC_022462(3) | NC_022463(3) | NC_023084(3) | |
| | | | | NC_024541(3) | NC_024543(3) | NC_024659(3) | |
| | | | | NC_024660(3) | NC_024661(3) | NC_024662(3) | |
| | | | | NC_024663(3) | NC_026197(3) | NC_026690(3) | |
| | | | | NC_026691(3) | NC_030180(3) | NC_030539(3) | |
| | | | | NC_030609(3) | NC_030784(3) | NC_030786(3) | |
| | | | | NC_030787(3) | NC_030789(3) | NC_031186(3) | |
| | | | | NC_031187(3) | NC_031428(3) | NC_031892(3) | |
| | | | | NC_033502(3) | NC_034331(3) | NC_034371(3) | |
| | | | | NC_034640(3) | NC_034641(3) | NC_034677(3) | |
| | | | | NC_034678(3) | NC_034913(3) | NC_034914(3) | |
| | | | | NC_034915(3) | NC_035418(3) | NC_033501(4) | |
| | | | | NC_019616(5) | NC_029704(5) | | |
| SRR6331521 | 3 | 4149 | family | NC_026202(3) | NC_031437(3) | | success |

Continued on next page

| SRA Accession | Depth | TaxID | Rank | Related References with Depth | Status |
|---|---|---|---|---|---|
| SRR6425047 | 5 | 2231393 | no rank | NC_009259(5) NC_013843(5) NC_016708(5)<br>NC_018051(5) NC_021091(5) NC_023090(5)<br>NC_025909(5) NC_027677(5) NC_029406(5)<br>NC_031429(5) NC_034742(5) NC_034743(5)<br>NC_034774(5) NC_002694(6) NC_003119(6)<br>NC_007942(6) NC_011163(6) NC_011828(6)<br>NC_014057(6) NC_014063(6) NC_021636(6)<br>NC_021645(6) NC_021646(6) NC_021647(6)<br>NC_021648(6) NC_021649(6) NC_021650(6)<br>NC_022868(6) NC_024034(6) NC_024035(6)<br>NC_024036(6) NC_024038(6) NC_024166(6)<br>NC_025743(6) NC_025744(6) NC_025745(6)<br>NC_027073(6) NC_027074(6) NC_027075(6)<br>NC_027076(6) NC_027077(6) NC_027078(6)<br>NC_027079(6) NC_027080(6) NC_027148(6)<br>NC_027149(6) NC_027150(6) NC_027151(6)<br>NC_027152(6) NC_027153(6) NC_027154(6)<br>NC_027155(6) NC_029828(6) NC_030329(6)<br>NC_032066(6) NC_032691(6) NC_034229(6)<br>NC_035228(6) NC_035229(6) NC_028171(7) | success |

Continued on next page

Table S7 – *Continued from previous page*

| SRA Accession | Depth | TaxID | Rank | Related References with Depth | Status |
|---|---|---|---|---|---|
| SRR6425048 | 5 | 2231393 | no rank | NC_009259(5) NC_013843(5) NC_016708(5)<br>NC_018051(5) NC_021091(5) NC_023090(5)<br>NC_025909(5) NC_027677(5) NC_029406(5)<br>NC_031429(5) NC_034742(5) NC_034743(5)<br>NC_034774(5) NC_002694(6) NC_003119(6)<br>NC_007942(6) NC_011163(6) NC_011828(6)<br>NC_014057(6) NC_014063(6) NC_021636(6)<br>NC_021645(6) NC_021646(6) NC_021647(6)<br>NC_021648(6) NC_021649(6) NC_021650(6)<br>NC_022868(6) NC_024034(6) NC_024035(6)<br>NC_024036(6) NC_024038(6) NC_024166(6)<br>NC_025743(6) NC_025744(6) NC_025745(6)<br>NC_027073(6) NC_027074(6) NC_027075(6)<br>NC_027076(6) NC_027077(6) NC_027078(6)<br>NC_027079(6) NC_027080(6) NC_027148(6)<br>NC_027149(6) NC_027150(6) NC_027151(6)<br>NC_027152(6) NC_027153(6) NC_027154(6)<br>NC_027155(6) NC_029828(6) NC_030329(6)<br>NC_032066(6) NC_032691(6) NC_034229(6)<br>NC_035228(6) NC_035229(6) NC_028171(7) | success |

Continued on next page

Table S7 – *Continued from previous page*

| SRA Accession | Depth | TaxID | Rank | Related References with Depth | Status |
|---|---|---|---|---|---|
| SRR6425049 | 5 | 2231393 | no rank | NC_009259(5) NC_013843(5) NC_016708(5) NC_018051(5) NC_021091(5) NC_023090(5) NC_025909(5) NC_027677(5) NC_029406(5) NC_031429(5) NC_034742(5) NC_034743(5) NC_034774(5) NC_002694(6) NC_003119(6) NC_007942(6) NC_011163(6) NC_011828(6) NC_014057(6) NC_014063(6) NC_021636(6) NC_021645(6) NC_021646(6) NC_021647(6) NC_021648(6) NC_021649(6) NC_021650(6) NC_022868(6) NC_024034(6) NC_024035(6) NC_024036(6) NC_024038(6) NC_024166(6) NC_025743(6) NC_025744(6) NC_025745(6) NC_027073(6) NC_027074(6) NC_027075(6) NC_027076(6) NC_027077(6) NC_027078(6) NC_027079(6) NC_027080(6) NC_027148(6) NC_027149(6) NC_027150(6) NC_027151(6) NC_027152(6) NC_027153(6) NC_027154(6) NC_027155(6) NC_029828(6) NC_030329(6) NC_032066(6) NC_032691(6) NC_034229(6) NC_035228(6) NC_035229(6) NC_028171(7) | success |
| SRR6425628 | 4 | 167487 | subfamily | NC_024292(3) NC_033354(3) NC_021423(4) | success |
| SRR6425646 | 4 | 167487 | subfamily | NC_024292(3) NC_033354(3) NC_021423(4) | success |
| SRR6425647 | 4 | 167487 | subfamily | NC_024292(3) NC_033354(3) NC_021423(4) | success |
| SRR6425648 | 4 | 167487 | subfamily | NC_024292(3) NC_033354(3) NC_021423(4) | success |
| SRR6425651 | 4 | 167483 | subfamily | NC_025657(3) NC_025655(4) NC_025656(4) NC_025658(4) | success |

Continued on next page

Table S7 – *Continued from previous page*

| SRA Accession | Depth | TaxID | Rank | Related References with Depth | Status |
|---|---|---|---|---|---|
| SRR6513658 | 3 | 3814 | subfamily | NC_009259(6) NC_013843(6) NC_016708(6) NC_018051(6) NC_021091(6) NC_023090(6) NC_025909(6) NC_027677(6) NC_029406(6) NC_031429(6) NC_034742(6) NC_034743(6) NC_034774(6) NC_002694(7) NC_003119(7) NC_007942(7) NC_011163(7) NC_011828(7) NC_014057(7) NC_014063(7) NC_021636(7) NC_021645(7) NC_021646(7) NC_021647(7) NC_021648(7) NC_021649(7) NC_021650(7) NC_022868(7) NC_024034(7) NC_024035(7) NC_024036(7) NC_024038(7) NC_024166(7) NC_025743(7) NC_025744(7) NC_025745(7) NC_027073(7) NC_027074(7) NC_027075(7) NC_027076(7) NC_027077(7) NC_027078(7) NC_027079(7) NC_027080(7) NC_027148(7) NC_027149(7) NC_027150(7) NC_027151(7) NC_027152(7) NC_027153(7) NC_027154(7) NC_027155(7) NC_029828(7) NC_030329(7) NC_032066(7) NC_032691(7) NC_034229(7) NC_035228(7) NC_035229(7) NC_028171(8) | success |

Continued on next page

Table S7 – *Continued from previous page*

| SRA Accession | Depth | TaxID | Rank | Related References with Depth | Status |
|---|---|---|---|---|---|
| SRR6821984 | 3 | 3700 | family | NC_000932(3) NC_009265(3) NC_009266(3)<br>NC_009267(3) NC_009268(3) NC_009269(3)<br>NC_009270(3) NC_009271(3) NC_009272(3)<br>NC_009273(3) NC_009274(3) NC_009275(3)<br>NC_016734(3) NC_018565(3) NC_021102(3)<br>NC_023367(3) NC_024469(3) NC_026445(3)<br>NC_026446(3) NC_027693(3) NC_028170(3)<br>NC_028272(3) NC_028415(3) NC_028517(3)<br>NC_028726(3) NC_028727(3) NC_028728(3)<br>NC_029253(3) NC_029254(3) NC_029331(3)<br>NC_029332(3) NC_029333(3) NC_029334(3)<br>NC_029335(3) NC_029336(3) NC_029337(3)<br>NC_029378(3) NC_029379(3) NC_030346(3)<br>NC_030347(3) NC_030348(3) NC_030349(3)<br>NC_030350(3) NC_030351(3) NC_030450(3)<br>NC_030515(3) NC_030516(3) NC_030775(3)<br>NC_033499(3) NC_033500(3) NC_034287(3)<br>NC_034299(3) NC_034357(3) NC_034359(3)<br>NC_034360(3) NC_034361(3) NC_034362(3)<br>NC_034363(3) NC_034365(3) NC_034366(3)<br>NC_034367(3) NC_035303(3) NC_034286(4)<br>NC_034379(4) | success |

Continued on next page

Table S7 – *Continued from previous page*

| SRA Accession | Depth | TaxID | Rank | Related References with Depth | Status |
|---|---|---|---|---|---|
| SRR6940055 | 3 | 4136 | family | NC_020098(3) NC_023102(3) NC_026291(3) NC_027262(3) NC_028533(3) NC_034693(3) NC_029370(4) NC_029817(4) NC_029818(4) NC_029819(4) NC_029820(4) NC_029821(4) NC_029822(4) NC_029823(4) NC_029824(4) NC_029825(4) NC_030755(4) NC_030756(4) NC_030757(4) NC_031433(4) NC_031434(4) NC_031874(4) NC_032054(4) NC_035143(4) NC_020431(5) NC_027259(5) NC_035233(5) | success |
| SRR6940081 | 3 | 4136 | family | NC_020098(3) NC_023102(3) NC_026291(3) NC_027262(3) NC_028533(3) NC_034693(3) NC_029370(4) NC_029817(4) NC_029818(4) NC_029819(4) NC_029820(4) NC_029821(4) NC_029822(4) NC_029823(4) NC_029824(4) NC_029825(4) NC_030755(4) NC_030756(4) NC_030757(4) NC_031433(4) NC_031434(4) NC_031874(4) NC_032054(4) NC_035143(4) NC_020431(5) NC_027259(5) NC_035233(5) | success |

Continued on next page

Table S7 – *Continued from previous page*

| SRA Accession | Depth | TaxID | Rank | Related References with Depth | Status |
|---|---|---|---|---|---|
| SRR7223704 | 3 | 3700 | family | NC_000932(3) NC_009265(3) NC_009266(3)<br>NC_009267(3) NC_009268(3) NC_009269(3)<br>NC_009270(3) NC_009271(3) NC_009272(3)<br>NC_009273(3) NC_009274(3) NC_009275(3)<br>NC_016734(3) NC_018565(3) NC_021102(3)<br>NC_023367(3) NC_024469(3) NC_026445(3)<br>NC_026446(3) NC_027693(3) NC_028170(3)<br>NC_028272(3) NC_028415(3) NC_028517(3)<br>NC_028726(3) NC_028727(3) NC_028728(3)<br>NC_029253(3) NC_029254(3) NC_029331(3)<br>NC_029332(3) NC_029333(3) NC_029334(3)<br>NC_029335(3) NC_029336(3) NC_029337(3)<br>NC_029378(3) NC_029379(3) NC_030346(3)<br>NC_030347(3) NC_030348(3) NC_030349(3)<br>NC_030350(3) NC_030351(3) NC_030450(3)<br>NC_030515(3) NC_030516(3) NC_030775(3)<br>NC_033499(3) NC_033500(3) NC_034287(3)<br>NC_034299(3) NC_034357(3) NC_034359(3)<br>NC_034360(3) NC_034361(3) NC_034362(3)<br>NC_034363(3) NC_034365(3) NC_034366(3)<br>NC_034367(3) NC_035303(3) NC_034286(4)<br>NC_034379(4) | success |

Continued on next page

Table S7 – *Continued from previous page*

| SRA Accession | Depth | TaxID | Rank | Related References with Depth | Status |
| --- | --- | --- | --- | --- | --- |
| SRR7223710 | 3 | 3700 | family | NC_000932(3) NC_009265(3) NC_009266(3)<br>NC_009267(3) NC_009268(3) NC_009269(3)<br>NC_009270(3) NC_009271(3) NC_009272(3)<br>NC_009273(3) NC_009274(3) NC_009275(3)<br>NC_016734(3) NC_018565(3) NC_021102(3)<br>NC_023367(3) NC_024469(3) NC_026445(3)<br>NC_026446(3) NC_027693(3) NC_028170(3)<br>NC_028272(3) NC_028415(3) NC_028517(3)<br>NC_028726(3) NC_028727(3) NC_028728(3)<br>NC_029253(3) NC_029254(3) NC_029331(3)<br>NC_029332(3) NC_029333(3) NC_029334(3)<br>NC_029335(3) NC_029336(3) NC_029337(3)<br>NC_029378(3) NC_029379(3) NC_030346(3)<br>NC_030347(3) NC_030348(3) NC_030349(3)<br>NC_030350(3) NC_030351(3) NC_030450(3)<br>NC_030515(3) NC_030516(3) NC_030775(3)<br>NC_033499(3) NC_033500(3) NC_034287(3)<br>NC_034299(3) NC_034357(3) NC_034359(3)<br>NC_034360(3) NC_034361(3) NC_034362(3)<br>NC_034363(3) NC_034365(3) NC_034366(3)<br>NC_034367(3) NC_035303(3) NC_034286(4)<br>NC_034379(4) | success |

Continued on next page

Table S7 – *Continued from previous page*

| SRA Accession | Depth | TaxID | Rank | Related References with Depth | Status |
|---|---|---|---|---|---|
| SRR8082590 | 3 | 3646 | order | NC_024060(3) NC_024061(3) NC_024062(3) | success |
| | | | | NC_024063(3) NC_024064(3) NC_024065(3) | |
| | | | | NC_024066(3) NC_024067(3) NC_026986(3) | |
| | | | | NC_030517(3) NC_030534(3) NC_030544(3) | |
| | | | | NC_030545(3) NC_030546(3) NC_030547(3) | |
| | | | | NC_030548(3) NC_030549(3) NC_030550(3) | |
| | | | | NC_030551(3) NC_030552(3) NC_030553(3) | |
| | | | | NC_030554(3) NC_030555(3) NC_030556(3) | |
| | | | | NC_030557(3) NC_030558(3) NC_030559(3) | |
| | | | | NC_030560(3) NC_030561(3) NC_030562(3) | |
| | | | | NC_030563(3) NC_030564(3) NC_030565(3) | |
| | | | | NC_030566(3) NC_030567(3) NC_030568(3) | |
| | | | | NC_030569(3) NC_030570(3) NC_030571(3) | |
| | | | | NC_030572(3) NC_030573(3) NC_030574(3) | |
| | | | | NC_030575(3) NC_030576(3) NC_030577(3) | |
| | | | | NC_030578(3) NC_030579(3) NC_030580(3) | |
| | | | | NC_030581(3) NC_030582(3) NC_030583(3) | |
| | | | | NC_030601(3) NC_034285(3) NC_008235(4) | |
| | | | | NC_009143(4) NC_024681(4) NC_024734(4) | |
| | | | | NC_024735(4) NC_024747(4) NC_026462(4) | |
| | | | | NC_026722(4) NC_027425(4) NC_028350(4) | |
| | | | | NC_028504(4) NC_031371(4) NC_031398(4) | |
| | | | | NC_032060(4) NC_032368(4) NC_032717(4) | |
| | | | | NC_033876(4) NC_010433(5) NC_012224(5) | |
| | | | | NC_015308(5) NC_016736(5) NC_034803(5) | |
| | | | | NC_033910(7) | |

Continued on next page

Table S7 – *Continued from previous page*

| SRA Accession | Depth | TaxID | Rank | Related References with Depth | Status |
|---|---|---|---|---|---|
| SRR8481357 | 5 | 2231393 | no rank | NC_009259(5) NC_013843(5) NC_016708(5) NC_018051(5) NC_021091(5) NC_023090(5) NC_025909(5) NC_027677(5) NC_029406(5) NC_031429(5) NC_034742(5) NC_034743(5) NC_034774(5) NC_002694(6) NC_003119(6) NC_007942(6) NC_011163(6) NC_011828(6) NC_014057(6) NC_014063(6) NC_021636(6) NC_021645(6) NC_021646(6) NC_021647(6) NC_021648(6) NC_021649(6) NC_021650(6) NC_022868(6) NC_024034(6) NC_024035(6) NC_024036(6) NC_024038(6) NC_024166(6) NC_025743(6) NC_025744(6) NC_025745(6) NC_027073(6) NC_027074(6) NC_027075(6) NC_027076(6) NC_027077(6) NC_027078(6) NC_027079(6) NC_027080(6) NC_027148(6) NC_027149(6) NC_027150(6) NC_027151(6) NC_027152(6) NC_027153(6) NC_027154(6) NC_027155(6) NC_029828(6) NC_030329(6) NC_032066(6) NC_032691(6) NC_034229(6) NC_035228(6) NC_035229(6) NC_028171(7) | success |

Continued on next page

Table S7 – *Continued from previous page*

| SRA Accession | Depth | TaxID | Rank | Related References with Depth | Status |
|---|---|---|---|---|---|
| SRR8481359 | 5 | 2231393 | no rank | NC_009259(5) NC_013843(5) NC_016708(5)<br>NC_018051(5) NC_021091(5) NC_023090(5)<br>NC_025909(5) NC_027677(5) NC_029406(5)<br>NC_031429(5) NC_034742(5) NC_034743(5)<br>NC_034774(5) NC_002694(6) NC_003119(6)<br>NC_007942(6) NC_011163(6) NC_011828(6)<br>NC_014057(6) NC_014063(6) NC_021636(6)<br>NC_021645(6) NC_021646(6) NC_021647(6)<br>NC_021648(6) NC_021649(6) NC_021650(6)<br>NC_022868(6) NC_024034(6) NC_024035(6)<br>NC_024036(6) NC_024038(6) NC_024166(6)<br>NC_025743(6) NC_025744(6) NC_025745(6)<br>NC_027073(6) NC_027074(6) NC_027075(6)<br>NC_027076(6) NC_027077(6) NC_027078(6)<br>NC_027079(6) NC_027080(6) NC_027148(6)<br>NC_027149(6) NC_027150(6) NC_027151(6)<br>NC_027152(6) NC_027153(6) NC_027154(6)<br>NC_027155(6) NC_029828(6) NC_030329(6)<br>NC_032066(6) NC_032691(6) NC_034229(6)<br>NC_035228(6) NC_035229(6) NC_028171(7) | success |

Continued on next page

Table S7 – *Continued from previous page*

| SRA Accession | Depth | TaxID | Rank | Related References with Depth | | | Status |
|---|---|---|---|---|---|---|---|
| SRR8784049 | 7 | 2231393 | no rank | NC_009259(5) | NC_013843(5) | NC_016708(5) | success |
| | | | | NC_018051(5) | NC_021091(5) | NC_023090(5) | |
| | | | | NC_025909(5) | NC_027677(5) | NC_029406(5) | |
| | | | | NC_031429(5) | NC_034742(5) | NC_034743(5) | |
| | | | | NC_034774(5) | NC_002694(6) | NC_003119(6) | |
| | | | | NC_007942(6) | NC_011163(6) | NC_011828(6) | |
| | | | | NC_014057(6) | NC_014063(6) | NC_021636(6) | |
| | | | | NC_021645(6) | NC_021646(6) | NC_021647(6) | |
| | | | | NC_021648(6) | NC_021649(6) | NC_021650(6) | |
| | | | | NC_022868(6) | NC_024034(6) | NC_024035(6) | |
| | | | | NC_024036(6) | NC_024038(6) | NC_024166(6) | |
| | | | | NC_025743(6) | NC_025744(6) | NC_025745(6) | |
| | | | | NC_027073(6) | NC_027074(6) | NC_027075(6) | |
| | | | | NC_027076(6) | NC_027077(6) | NC_027078(6) | |
| | | | | NC_027079(6) | NC_027080(6) | NC_027148(6) | |
| | | | | NC_027149(6) | NC_027150(6) | NC_027151(6) | |
| | | | | NC_027152(6) | NC_027153(6) | NC_027154(6) | |
| | | | | NC_027155(6) | NC_029828(6) | NC_030329(6) | |
| | | | | NC_032066(6) | NC_032691(6) | NC_034229(6) | |
| | | | | NC_035228(6) | NC_035229(6) | NC_028171(7) | |

Continued on next page

Table S7 – *Continued from previous page*

| SRA Accession | Depth | TaxID | Rank | Related References with Depth | Status |
|---|---|---|---|---|---|
| SRR8784093 | 5 | 2231393 | no rank | NC_009259(5) NC_013843(5) NC_016708(5)<br>NC_018051(5) NC_021091(5) NC_023090(5)<br>NC_025909(5) NC_027677(5) NC_029406(5)<br>NC_031429(5) NC_034742(5) NC_034743(5)<br>NC_034774(5) NC_002694(6) NC_003119(6)<br>NC_007942(6) NC_011163(6) NC_011828(6)<br>NC_014057(6) NC_014063(6) NC_021636(6)<br>NC_021645(6) NC_021646(6) NC_021647(6)<br>NC_021648(6) NC_021649(6) NC_021650(6)<br>NC_022868(6) NC_024034(6) NC_024035(6)<br>NC_024036(6) NC_024038(6) NC_024166(6)<br>NC_025743(6) NC_025744(6) NC_025745(6)<br>NC_027073(6) NC_027074(6) NC_027075(6)<br>NC_027076(6) NC_027077(6) NC_027078(6)<br>NC_027079(6) NC_027080(6) NC_027148(6)<br>NC_027149(6) NC_027150(6) NC_027151(6)<br>NC_027152(6) NC_027153(6) NC_027154(6)<br>NC_027155(6) NC_029828(6) NC_030329(6)<br>NC_032066(6) NC_032691(6) NC_034229(6)<br>NC_035228(6) NC_035229(6) NC_028171(7) | success |
